# Foldamers Reveal and Validate Novel Therapeutic Targets Associated with Toxic α-Synuclein Self-Assembly

**DOI:** 10.1101/2021.05.08.443146

**Authors:** Jemil Ahmed, Tessa C. Fitch, Courtney M. Donnelly, Johnson A. Joseph, Mikaela M. Bassil, Ahyun Son, Chen Zhang, Aurélie Ledreux, Scott Horowitz, Yan Qin, Daniel Paredes, Sunil Kumar

## Abstract

Parkinson’s disease (PD) is a progressive neurodegenerative disorder for which there is no successful prevention or intervention. The pathological hallmark for PD involves the self-assembly of functional Alpha-Synuclein (αS) into non-functional amyloid structures. One of the potential therapeutic interventions against PD is the effective inhibition of αS aggregation. However, the bottleneck towards achieving this goal is the identification of αS domains/sequences that are essential for aggregation. Using a protein mimetic approach, we have identified αS sequences-based novel targets that are essential for aggregation and will have significant therapeutic implications. An extensive array of *in vitro, ex vivo*, and *in vivo* assays was utilized to validate αS sequences and their structural characteristics that are essential for aggregation and propagation of PD phenotypes. The study aids in developing significant mechanistic and therapeutic insights into various facets of αS aggregation, which will pave the way for novel and effective treatments for PD.

## INTRODUCTION

Alpha-Synuclein (αS) is a neuronal protein expressed at high levels in dopaminergic neurons and it is believed to be implicated in the regulation of synaptic vesicle trafficking and recycling, and neurotransmitter release^1-8^. The misfolding of αS leads to its self-aggregation, which is a pathological hallmark of PD^1-8^. Therefore, modulation of αS aggregation is a promising therapeutic intervention for PD^1-10^. The identification and specific targeting of sequences or domains that initiate αS aggregation could promise potent antagonism of the αS self-assembly. A few small molecules have been shown to inhibit αS aggregation (11 and ref. within), however, limited atomic-level understanding is available of the ligand-αS interaction, which restricted the further optimization of the antagonists against αS aggregation. More importantly, limited progress has been made in the identification of factors that are associated with αS aggregation, e.g. αS sequences that initiate aggregation. Mutation studies enable the identification of αS sequences/domains that are important for aggregation^12-16^. However, no study has been directed to validate these αS sequences as novel targets. Here, we have utilized a foldamer-based approach in tandem with a mutation study that allowed the identification and validation of novel αS sequences as key therapeutic targets that are essential for the initiation of αS aggregation.

Foldamers are dynamic ligands with the ability to mimic the topography and the chemical space of the secondary structure of proteins^17-22^. The diversity of chemical space can be conveniently tuned in foldamers, an essential property for the optimization of interactions with targets. Oligoquinoline (OQ)-based foldamers have been shown to modulate the self-assembly of islet amyloid polypeptide^22-26^ and Aβ peptide^21^, whose aggregation is associated with type 2 diabetes (T2D) and Alzheimer’s disease (AD), respectively.

We have utilized OQs to gain mechanistic and therapeutic insights into αS aggregation. Using an array of biophysical, cellular, *in vivo* assays and mutation studies, we have identified SK-129, a potent antagonist of αS aggregation in both *in vitro* and *in vivo* PD models. A 2D-NMR-based atomic-level investigation enabled the identification of the binding sites of SK-129 on αS, which were validated using fluorescent polarization and mutation studies. More importantly, we identified αS sequences as novel targets that are essential for the initiation of the aggregation. We also validated novel αS sequences by targeting them with OQs and rescued PD phenotypic readouts in cellular, neuronal, and *in vivo* PD models. SK-129 was a potent antagonist of the prion-like spread of αS seeds. The activity of SK-129 against the prion-like spread of αS seeds was confirmed using distinct αS seed polymorphs generated from the recombinant αS and extracted from the substantia nigra of the post mortem brain of PD patient. The activity of SK-129 was also confirmed in a novel intracellular assay for the prion-like spread of αS seeds. Overall, SK-129 interacts at the N-terminus of αS monomer, induces or stabilizes a helical conformation, and modulates both *de novo* aggregation and the prion-like spread of αS seeds. We used a chemical tool to identify and validate novel αS sequences with structural insights that are essential for the aggregation and associated with PD phenotypes. The study will have significant mechanistic and therapeutic implications, which will aid in expediting treatments for PD.

## RESULTS

### Biophysical characterization of foldamers with αS

The OQs with carboxylic acid and hydrophobic side chains have been shown to modulate the self-assembly of amyloid proteins by specifically targeting sequences that are rich in positively charged and hydrophobic side chain residues^21-26^. Therefore, we utilized an established library of OQs with carboxylic acid and various hydrophobic groups as side chains (Fig. 1a-c). The library was screened against αS aggregation using Thioflavin T (ThT) dye-based amyloid assay^27^. The aggregation kinetics of 100 μM αS (in 1 × PBS buffer) was characterized by a sigmoidal curve with a t50 (time to reach 50% fluorescence) of ∼38.1±1.8 h. The screening led to the identification of SK-129 as the most potent antagonist of wild type (WT) αS (and αS mutants, αSA30P and αSA53T) aggregation at equimolar and sub-stoichiometric ratios (Fig. 1c,d and Supplementary Fig. 1). SK-129 inhibits αS aggregation under *de novo* and lipid membrane conditions at an equimolar ratio (Supplementary Fig. 2a), which was also validated by transmission electron microscopy (TEM) images (Supplementary Fig. 2b,c).

**Fig. 1.**
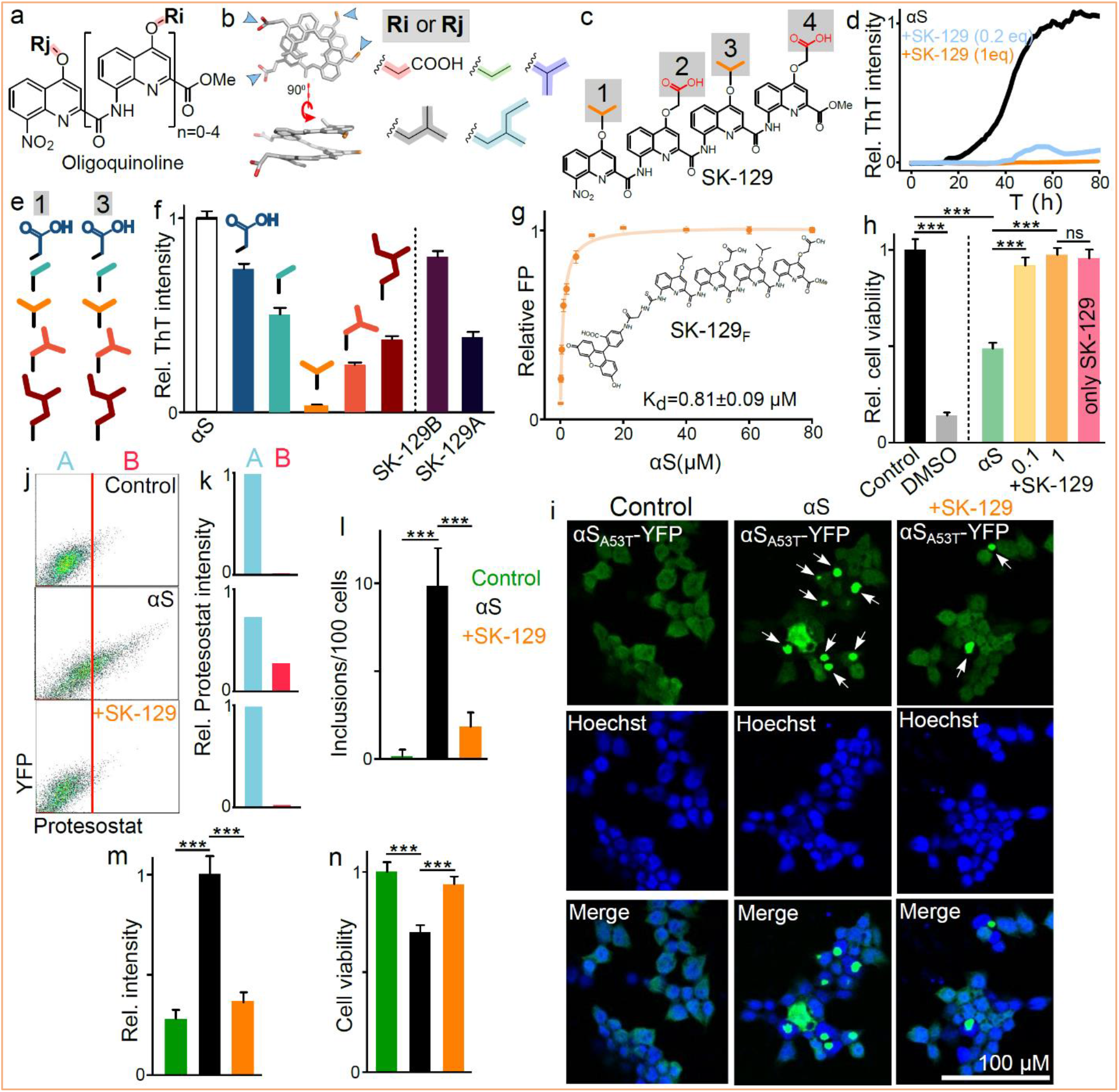
Characterization of the antagonist activity of SK-129 against αS aggregation. The chemical (**a**) and crystal (**b**) structures of OQs and surface functionalities. **c**, Chemical structure of SK-129. **d**, The aggregation profile of 100 µM αS in the absence and presence of SK-129 at the indicated molar ratios. **e**, The chemical structures and antagonist activities (**f**) of 100 μM SK-129 derivatives against 100 μM αS aggregation. **g**, The FP titration curve to determine the binding affinity between 10 μM SK-129F and αS. **h**, The statistical analysis of the relative viability of SH-SY5Y cells in the presence of 10 μM αS and the αS-SK-129 complex at the indicated molar ratios. **i**, Confocal images of HEK cells treated with the indicated conditions. Inclusions=white arrows, Hoechst (blue), merge = Hoechst and αSA53T-YFP. **j**, The Flow cytometry analysis of HEK cells treated with the indicated conditions. The x-axis represents αSA53T-YFP aggregates containing cells stained with ProteoStat dye. **k**, A and B represent relative % of HEK cells without and with αSA53T-YFP aggregates, respectively. **l**, The number of inclusions in HEK cells observed for the indicated conditions. The relative intensity of ProteoStat-stained aggregates (**m**) and relative viability (**n**) of HEK cells under the indicated conditions. Statistical significance, one-way analysis of variance (ANOVA) with Tukey’s multiple comparison test. ^*^p<0.05, ^**^p<0.01, ^***^p<0.001.

The antagonist activity of SK-129 is predominantly a consequence of the side chains. Among analogs (with varying hydrophobicity), SK-129 was the most potent antagonist of αS aggregation (Fig. 1e,f and Supplementary Scheme 3), which indicates that moderate hydrophobicity at positions 1 and 3 is required to achieve the optimal activity. The positioning of the side chains was important for SK-129’s activity as scrambling of side chains led to significantly diminished activity (Fig. 1f, Supplementary Scheme 1,2). The antagonist activity of SK-129 was much better than Epigallocatechin Gallate (EGCG), a potent antagonist of αS aggregation^28^. Gel shift assay shows that SK-129 potently inhibits αS aggregation; however, higher-order toxic oligomers (n>5)^29-30^ were observed in the presence of EGCG (Supplementary Fig. 3). We next determine the binding affinity of SK-129 using fluorescence polarization (FP) titration between a fluorescent analog of SK-129 (SK-129F) (Supplementary Scheme 3) and αS, which yielded a Kd of 0.81±0.09 μM (Fig. 1g) and a binding stoichiometry of 1:1 (αS:SK-129F) (Supplementary Fig. 4). The complex of SK-129F-αS was used for a displacement titration with SK-129, which yielded a Kd of 0.72±0.06 μM (Supplementary Fig. 5). More importantly, SK-129F (SK-129F-αS complex) could be used as a novel tool for a high throughput assay to screen and identify high-affinity ligands for αS. Also, SK-129 is a very specific antagonist of αS aggregation. The Kd of SK-129 for Aβ42 was more than 15-fold higher than αS (Supplementary Fig. 6a). Corroborating this specificity, SK-129 did not show any noticeable effect on Aβ aggregation (Supplementary Fig. 6b).

The aggregation of αS is associated with toxicity^1-8^; therefore, we tested the efficacy of SK-129 on the neuroblastoma SH-SY5Y cell line. The aggregation of αS was carried out in the absence and presence of SK-129 at an equimolar ratio, and the solutions were tested in SH-SY5Y^31^. The viability of SH-SY5Y cells decreased to 48±3% upon exposure to 10 μM αS for 24 h; which was rescued to 91±4% and 97±4% in the presence of SK-129 at molar ratios of 1:0.1 and 1:1 (αS:SK-129), respectively (Fig. 1h). The rescue of toxicity by SK-129 in SH-SY5Y cells was similar at higher concentrations of αS (25 μM and 50 μM) (Supplementary Fig. 7). To confirm that SK-129 did not generate seed-competent structures during the inhibition of αS aggregation, we utilized two HEK cell lines, which stably express YFP-labeled WT αS (αS-YFP) and a familial mutant, A53T (αS-A53T-YFP)^32-33^. Both HEK cells have been shown to template endogenous monomeric αS-A53T-YFP (αS-YFP) into fibers when transfected with αS fibers with lipofectamine 3000 (Fig. 1i)^32-33^, which is detected by intracellular fluorescent puncta (Fig. 1i, white arrows). αS was aggregated in the absence and presence of SK-129 at an equimolar ratio and the HEK cells were treated with αS fibers (7 μM in monomeric αS) for 24 h (Fig. 1i). A very small number of inclusions (αS-A53T-YFP or αS-YFP) were observed in the presence of SK-129 (Fig. 1i). The inclusions (αS-A53T-YFP) were quantified by confocal microscopy (Fig. 1i,l), flow cytometry (inclusions stained with ProteoStat dye^34^, Fig. 1j,k), and a 96-well plate reader (using ProteoStat dye, Fig. 1m), which were alleviated significantly in the presence of SK-129. The viability of HEK cells improved from 68% to 94% (αS aggregated solution in the presence of SK-129), which was determined using the (3-(4,5-dimethylthiazol-2-yl)-2,5-diphenyltetrazolium bromide) (MTT) reduction assay (Fig. 1n). SK-129 was equally effective in inhibiting aggregation and cytotoxicity in HEK cells that were expressing intracellular αS-YFP (Supplementary Fig. 8).

### Identification of the binding site of SK-129 on αS

The N-terminal domain spanning residues 1-90 plays a significant role in αS aggregation^8,12,35-38^. Therefore, we hypothesized that SK-129 could be interacting with the N-terminus of αS for the potent inhibition. We utilized two-dimensional heteronuclear single quantum coherence NMR spectroscopy (2D NMR HSQC) for atomic-level insight into the binding site of SK-129 on αS. We collected the HSQC NMR of 70 μM ^15^N-^1^H-uniformly labeled αS in the absence and presence of SK-129 at an equimolar ratio (Fig. 2a-e) and compared the signal intensity of the amide peaks. The total changes in the intensity in the presence of SK-129 suggest that the binding site of SK-129 is toward the N-terminus of αS (Fig. 2a-e), more specifically SK-129 interacts and changes the conformation of four αS sequences, including 6-12, 15-23, 36-45, and 48-53 (Fig. 2a-e). The binding sites of SK-129 on αS contain lysine and hydrophobic residues; therefore, we propose that the carboxylic acid and the propyl side chains of SK-129 are involved in binding interactions with lysine and hydrophobic residues of αS.

**Fig. 2.**
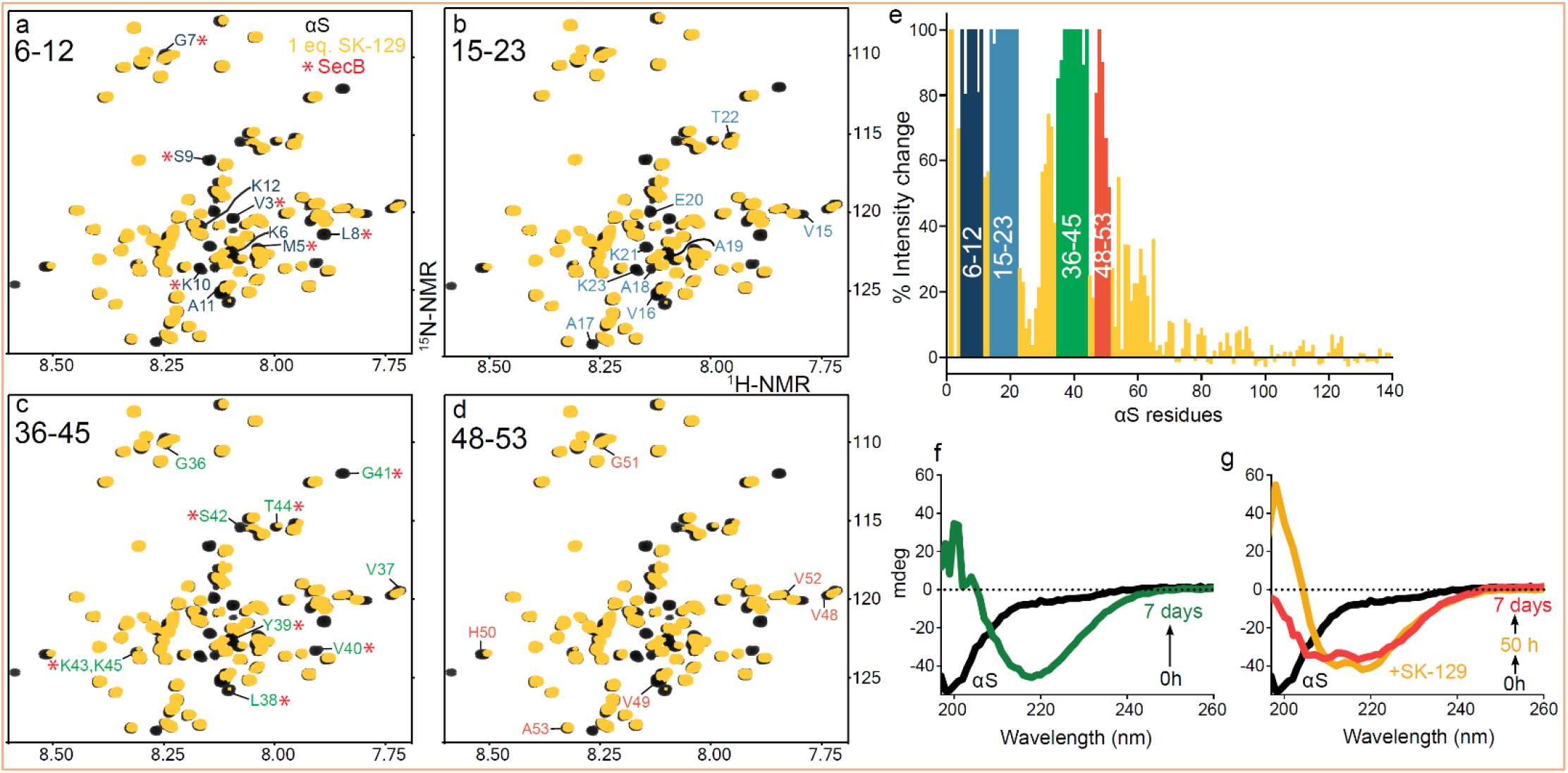
Structural characterization of the binding interaction between SK-129 and αS. **a-d**, Overlay of two-dimensional HSQC (^1^H, ^15^N) NMR spectra of 70 µM uniformly ^15^N-labelled αS in the absence (black) and presence (yellow) of SK-129 at an equimolar ratio. The largest attenuation in the volume of the backbone amide residue NMR signals are highlighted and assigned, which includes αS segments from 6-12 (**a**), 15-23 (**b**), 36-45 (**c**), and 48-53 (**d**). The change in the volume of amide backbone residue peaks of αS was compared between SK-129 and a molecular chaperone SecB (*, red) and the pronounced changes were observed in segments 6-12 (**a**) and 36-45 (**b**). **e**, Graphical presentation of the changes in the chemical shifts of the backbone amide residue peaks of ^15^N-labelled αS (70 µM) in the presence of SK-129 at an equimolar ratio. The colored sequences are the potential binding sites of SK-129 on αS. CD-based characterization of the aggregation kinetics of 35 µM αS in the absence (**f**) and presence (**g**) of SK-129 at an equimolar ratio. The spectra were recorded for 7 days.

### Effect of SK-129 on αS conformation

The NMR study also suggests that SK-129 induces α-helical conformation in αS. The intensity changes of αS residues in the presence of large unilamellar vesicles [0.875 mM, LUVs, 100 nm, DOPS, 1,2-dioleoyl-sn-glycero-3-phospho-L-serine (sodium salt)] were similar to those influenced by SK-129 at a higher molar ratio (1:2, αS:SK-129) (Supplementary Fig. 9-11). As αS samples α-helical conformations in the presence of LUVs^37-39^, we postulated that αS forms an α-helical conformation in the presence of SK-129. We utilized circular dichroism (CD) to study the interaction of αS with SK-129. The conformation of 35 μM αS transitioned from random coil to β-sheet in 7 days (Fig. 2f); however, the conformation of αS switched from random coil to α-helix and stayed in the same conformation in the presence of SK-129 at an equimolar ratio(Fig. 2g). We posit that the antagonist activity of SK-129 against αS aggregation is a consequence of the direct interaction with the N-terminus and the induction or stabilization of an α-helical conformation in αS. SK-129 behaved similarly under lipid catalyzed αS aggregation. SK-129 was a potent antagonist of LUVs catalyzed αS aggregation (Supplementary Fig. 2a. The CD spectra of 30 μM αS switched from an α-helix to a β-sheet conformation via an α-helical conformation in the presence of LUVs (375 μM, 100 nm, DOPS, Supplementary Fig. 12a)^37,39,40^. In marked contrast, αS remained in an α-helix conformation in the presence of LUVs and SK-129 at an equimolar ratio (Supplementary Fig. 12b). However, there was a decrease in the CD intensity of α-helix upon the addition of SK-129, which suggests that SK-129 might be competing against the lipid membrane for αS (Supplementary Fig. 12b). We surmise that the inhibition of the membrane-catalyzed oligomerization/aggregation of αS might be a consequence of the competition of αS between LUVs and SK-129.

We utilized HSQC NMR to monitor the effect of SK-129 on LUVs associated αS. SK-129 (70 μM) was added to the complex of ^15^N αS:LUVs (70 μM: 875 µM) and the intensity changes were compared with the αS:SK-129 complex (Supplementary Fig. 13). The NMR spectrum was a combination of αS-SK-129 and αS-LUVs complexes, which suggests an interchange of αS between SK-129 and LUVs (Supplementary Fig. 13). Collectively, our results suggest that SK-129 is inhibiting αS aggregation by partially competing with LUVs via transient interactions with αS and without completely displacing αS from LUVs.

To further confirm the binding sites of SK-129 on αS, we carried out a mutation study by systematically removing residues 6-12, 15-23, 36-45, or 48-53 from WT αS denoted as αS1, αS2, αS3, and αS4, respectively (Fig. 3a and Supplementary Fig. 14-15). The FP-based binding affinity of SK-129F for αS1 and αS2 mutants was 3-4 fold weaker (than WT αS) and very weak for both αS3 (∼8 fold) and αS4 mutants (>10 fold, Fig. 3b-e). We posit that SK-129 has multiple binding sites on αS with varying binding affinities or that the main binding site spans residues 36-53, and the intensity change of residues 6-12 and 15-23 is a consequence of the conformational switch in αS.

**Fig. 3.**
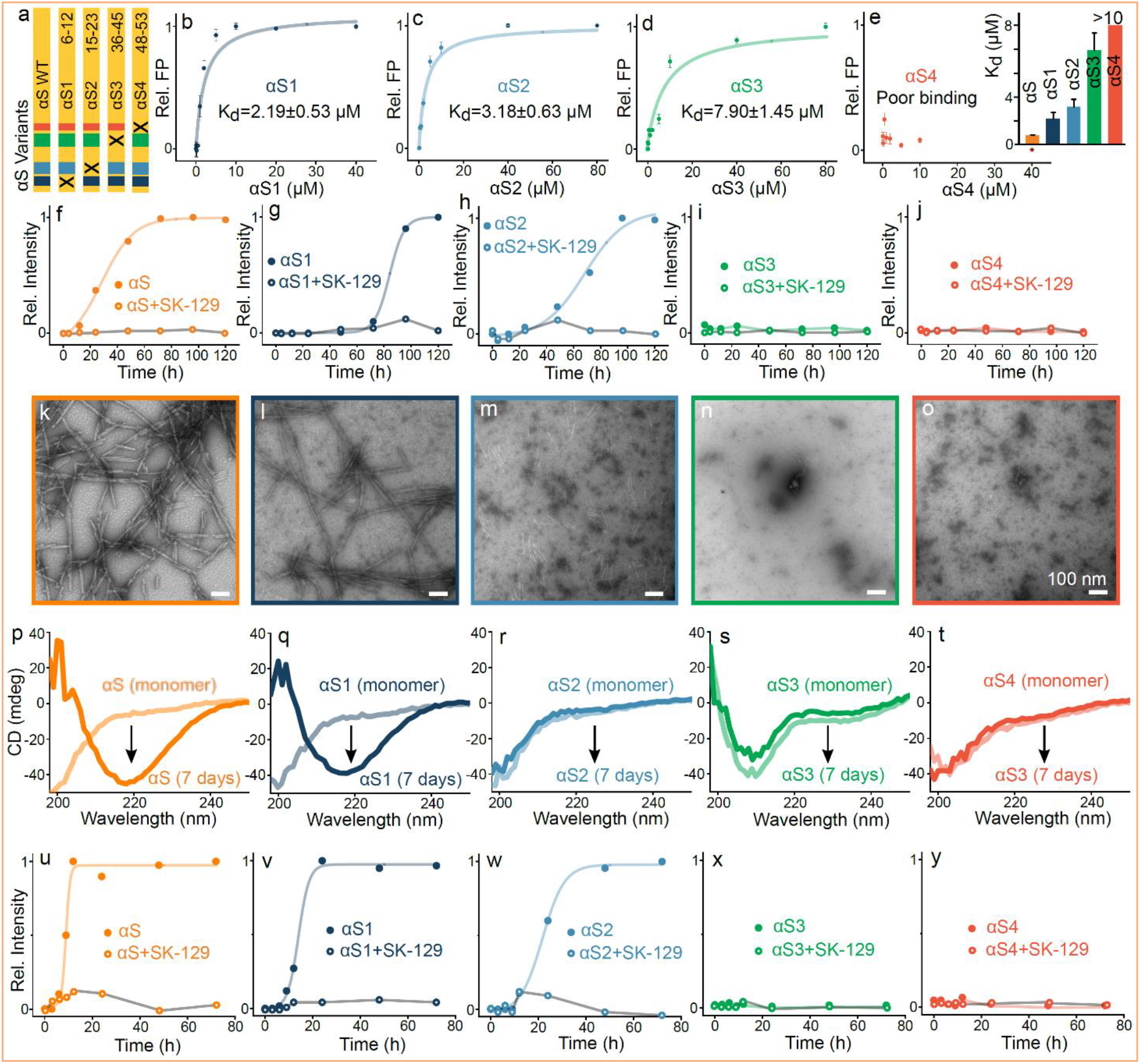
SK-129-based mutation study to delineate the role of αS sequences in aggregation. **a**, A schematic of the design of αS variants where “x” represents the deleted sequence from the WT αS. **b-e**, The fits for the FP titrations to determine the binding affinities between 10 µM SK-129_F_ and αS variants (inset). **f-j**, ThT fluorescence-based aggregation kinetic profiles of 100 µM αS variants in the absence (closed circle) and presence (open circle) of SK-129 at an equimolar ratio. The circles represent the average ThT intensity of three different experiments. **k-o**, TEM images of αS variants (100 µM) after aggregated them for seven days. **p-t**, CD spectra of monomeric (light color) and the aggregated (dark color) states of αS variants (35 µM). The same aggregated samples of αS variants were used for both CD and TEM images. **u-y**, Aggregation profiles of 100 µM αS variants catalyzed by preformed fibers of WT αS (10 µM in monomeric unit) in the absence (close circle) and presence (open circle) of SK-129 at an equimolar ratio. The circles represent the average ThT intensity of three different experiments. The fluorescence polarization titrations between SK129 and various proteins were conducted at least three times and each point in titrations represents the average of three data points. The reported error bars are the s.d.’s for multiple sets of experiments conducted on separate occasions.

### αS sequences essential for *de novo* and seed-catalyzed aggregation

SK-129 inhibits aggregation by interacting with four αS sequences; therefore, we hypothesize that these sequences might be essential to initiate αS aggregation. Therefore, we investigated the effect of these sequences on αS aggregation. Mutants αS3 and αS4 did not aggregate under our conditions via ThT and TEM (Fig. 3a,i,j,n,o and Supplementary Fig. 16), and their CD spectra were random coil (Fig. 3s,t). Mutants αS1 and αS2 aggregated with t50’s 3-4 fold higher than WT αS (Fig. 3a,f,g and Supplementary Fig. 16). The morphology of αS1 fibers was similar to WT αS (Fig. 3l); however, αS2 fibers were amorphous (Fig. 3l). Both WT αS and αS1 sampled β-sheet conformation (Fig. 3p,q); however, αS2 did not have the characteristics of a β-sheet conformation (Fig. 3r).

We also investigated the role of these αS sequences on the seed-catalyzed aggregation of αS. The WT αS seeds (10% monomer concentration) accelerated 100 μM αS aggregation by decreasing the t50 of WT αS (28.3±2.2 h), αS1 (84.1±3.6 h), αS2 (72.1±3.4 h) to WT αS (8.9±0.2 h), αS1 (14.0±0.6 h), αS2 (21.6±0.7 h) (Fig. 3u-y and Supplementary Fig. 17). The αS seeds did not template and aggregated mutants αS3 and αS4 (Fig. 3x,y and Supplementary Fig. 17), which suggests that the deleted sequences in αS3 and αS4 might be involved in seed catalyzed aggregation. SK-129 wholly suppressed the seed-catalyzed aggregation of WT αS, αS1 and αS2 (Fig. 3u-w and Supplementary Fig. 17) at an equimolar ratio. These experiments show that the sequences affected by SK-129 are important for αS aggregation.

The seed-catalyzed aggregation of αS conceptually mimics the prion-like spread of αS seeds, a phenomenon in which αS seeds enable the rapid conversion of functional and soluble αS into insoluble fibrils ^41-48^. An *in vitro* model of the prion-like spread of αS has been recapitulated in the protein misfolding cyclic amplification (PMCA) technique ^49-51^. In the PMCA assay, αS fibers are amplified for five cycles using αS monomer and seeds from the previous cycle. Additionally, αS seed polymorphs from different sources differ in mediating PD phenotypes, which is a consequence of the prion-like spread through infection pathways^51-53^. Therefore, we utilized two αS seed polymorphs, including recombinant αS seeds and αS seeds extracted from the substantia nigra of a PD brain and a control brain (post mortem condition) and used them in the PMCA assay (Fig. 4a). The Lewy bodies (LBs)/αS aggregates in the substantia nigra of the PD brain were confirmed using immunostaining (bluish/black, black arrows) and TEM (Fig. 4c,e). We did not observe LBs in the control brain (Fig. 4b,d). The PMCA assay sample (cycle 5) of the control brain extract was neither effective in seeding nor PK (proteinase K) resistant (Fig. 4f). However, the sample (cycle 5) from the PD brain was effective in seeding and was PK resistant (Fig. 4f). The sample was also effective at templating αS-_A53T_-YFP monomer into inclusions in HEK cells (Fig. 4i,k). The inclusions and toxicity increased gradually up to four days in the presence of the seeds from PD brain sample (Fig. 4i,k,l). Under the biological condition used, the control HEK cells were healthy up to 4 days and therefore, we decide to restrict our study up to four days.

**Fig. 4.**
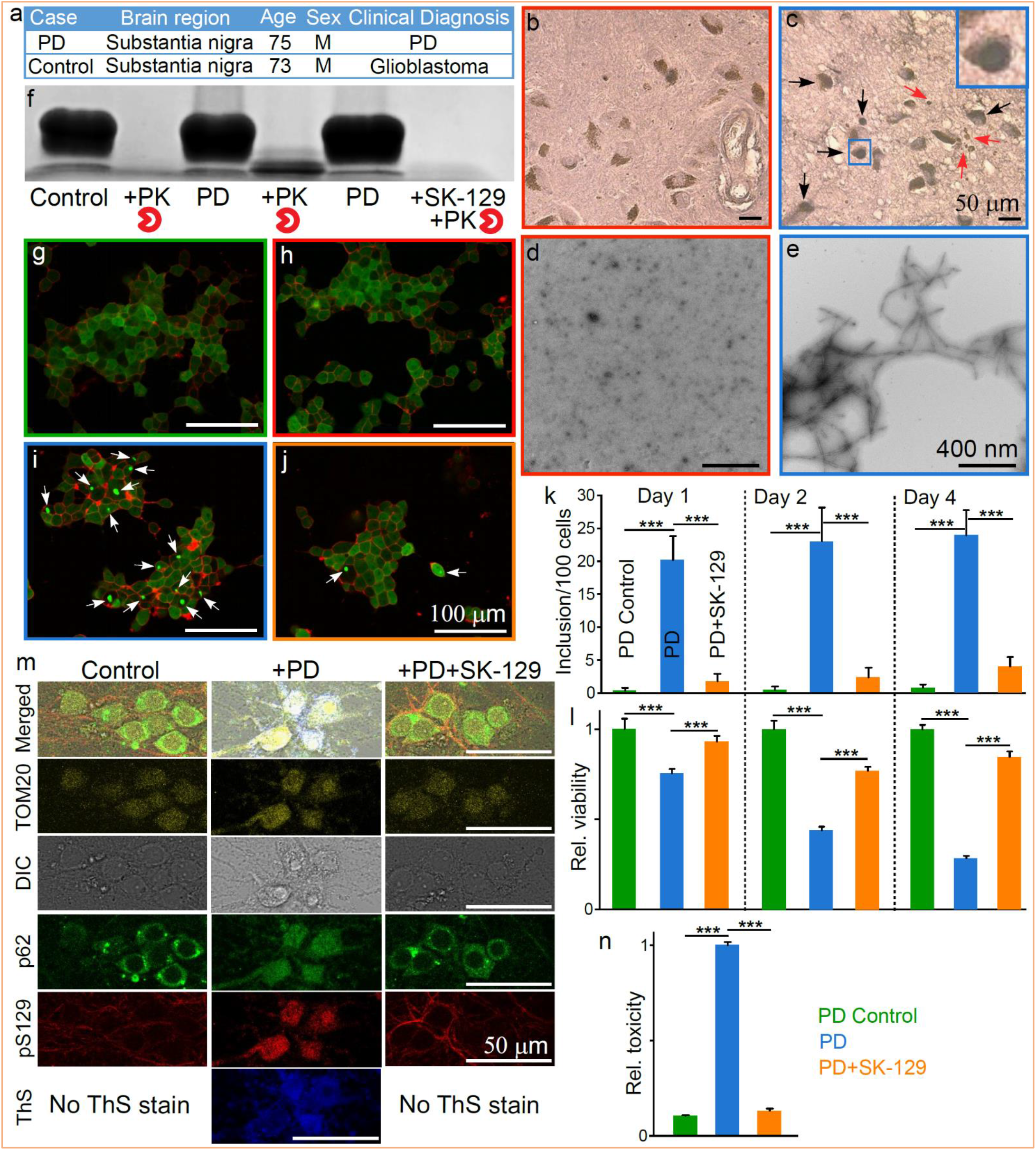
The assessment of the antagonist activity of SK-129 in *ex vivo* PD models **a**, The demographic and clinical information of the human brain tissues. Neuromelanin (brown) and αS immunostaining (LBs, bluish/black, black arrows, inset) in substantia nigra neurons from control (**b**) and PD (**c**) post mortem brain. Degenerating neurons and the extracellular neuromelanin debris from dying neurons (red arrows) were also visible. The hollow spaces in the PD brain demarcate cell loss. TEM images of the αS seeds extracted from the control (**d**) and PD brains (**e**). **f**, The αS stained western blot of the PMCA sample from the fifth cycle of the control and PD brain extracts after treatment with PK in the absence and presence of SK-129. Confocal images of HEK cells after treatment with control (**g**), control (**h**) and PD (**i**) brain extracts from PMCA sample (fifth cycle) and in the presence of SK-129 (**j**) at an equimolar ratio (**j**). **k, l**, Statistical analysis of the number of inclusions (**k**) and relative viability of HEK cells in the presence of PMCA samples (fifth cycle) from PD brain extracts under the indicated conditions. **m**, Confocal imaging of primary neurons treated with PMCA samples (fifth cycle) of control and PD brain extracts in the absence and presence of SK-129 at an equimolar ratio for 21 days. The primary neurons were stained with various markers, including LB biomarkers (pS129 and p62), mitochondria marker (TOM20), and ThS dye. **n**, Under matched conditions (to **m**), the neurotoxicity of primary neurons was measured using the LDH assay. *p<0.05, **p<0.01, ***p<0.001.

### Effect of SK-129 on the prion-like spread of αS seeds in an *ex vivo* PD model

We next investigated the effect of SK-129 at preventing seed-catalyzed aggregation from PD brain samples. The sample (cycle 5) of PD brain extract in the presence of SK-129 was neither aggregated nor PK resistant (Fig. 4a). Also, we observed a lower number of inclusions and improved cell viability for up to four days (Fig. 4j-l). We observed similar behavior of the PMCA sample (cycle 5) from recombinant αS seeds in the absence and presence of SK-129 at an equimolar ratio (Supplementary Fig. 18). The PMCA assay for recombinant protein leads to an abundance of αS fibers (Cycle 5), confirmed with high ThT signal, TEM image, PK resistance, inclusions and toxicity in HEK cells (Supplementary Fig. 18). In contrast, there was no formation of αS fibers for the PMCA assay (Cycle 5) in the presence of SK-129 as confirmed by ThT intensity, TEM image, PK cleavage, very low number of inclusions and rescue of toxicity in HEK cells (Supplementary Fig. 18).

To further confirm the effect of SK-129 on the formation of LBs and on the prion-like spread of αS seeds in the presence of the PD brain extract, we employed a more physiologically relevant model based on the primary rat hippocampal neurons. Using primary hippocampal neurons, an αS aggregation-based seeding model has been recently developed that recapitulates the key events of aggregation, seeding, and maturation of inclusions that mimic the bona fide LBs^48^. We incubated PMCA samples (cycle 5) of brain extracts in the absence and presence of SK-129 in primary hippocampal neurons for 21 days, a reported timeline to form matured LBs^48^. The neurons treated with PD sample (Fig. 4m) were stained positive for LBs biomarkers, including αS-pS-129 (phosphorylated residue 129) (red color) and ThS, a dye that specifically binds protein aggregates (blue color, Fig. 4m)^54^, p62 (autophagosome vesicles) and TOM20 (mitochondria) (Fig. 4m)^48^. The data suggest that αS inclusions recruit and sequester various organelles, proteins, and membranous structures^45,48,55-56^. We did not observe any colocalization of αS-pS-129 with p62 and TOM20 when neurons were treated with PD sample in the presence of SK-129 (Fig. 4m, +PD+SK-129). Also, no staining of αS-pS-129 was observed with ThS (Fig. 4m). Using lactate dehydrogenase (LDH) release assay, we observed very high neurotoxicity (∼8-fold higher than control) in primary neurons in the presence of PD sample; however, no toxicity was observed in the presence of PD sample treated with SK-129 (Fig. 4n).

### Effect of SK-129 on αS aggregation mediated PD phenotypes in an *in vivo* model

The antagonist activity of SK-129 against αS aggregation was tested *in vivo* using a *C elegans*-based PD model (NL5901). The ability of SK-129 to efficiently permeate cell membranes was confirmed by the parallel artificial membrane permeation assay (Fig. 5a) and confocal microscopy (using SK-129F, Fig. 5b). The NL5901 strain expresses WT αS-YFP in the body wall muscle cells^49,57^ and PD phenotypic readouts include a gradual increase in inclusions (αS-YFP) in body wall muscle cells and a decline in motility during aging (Fig. 5c,e,f)^49,57^. The NL5901 strain was treated with 15 μM SK-129 at the larval stage and incubated with and without SK-129 for nine days. The inclusions (αS-YFP) were counted manually using confocal microscopy. We observed a high number of inclusions (∼33 inclusions/*C elegans*) (Fig. 5c,e and Movie S1); however, there was a substantial decline in inclusions in the presence of SK-129 (∼8-9 inclusions/*C elegans*) (Fig. 5d,e and Movie S2), suggesting that SK-129 permeates the body wall muscle cell membrane and inhibits αS aggregation (Fig. 5d,e). The motility rate of the NL5901 strain decreases during the aging process as a consequence of αS inclusions. We utilized a newly developed WMicroTracker ARENA plate reader to measure the locomotion (overall activity counts) of NL5901 in the absence and presence of SK-129^58,59^. The overall activity of NL5901 displayed a gradual decline in the activity in comparison to the WT model of *C elegans* (N2) (Fig. 5f and Supplementary Fig. 19); however, NL5901 treated with 15 µM SK-129 at the larval stage resulted in a significant improvement in the overall activity (Fig. 5f and Supplementary Fig. 19). The overall activity of NL590 treated with SK-129 was closer to N2 (Fig. 5f and Supplementary Fig. 19).

**Fig. 5.**
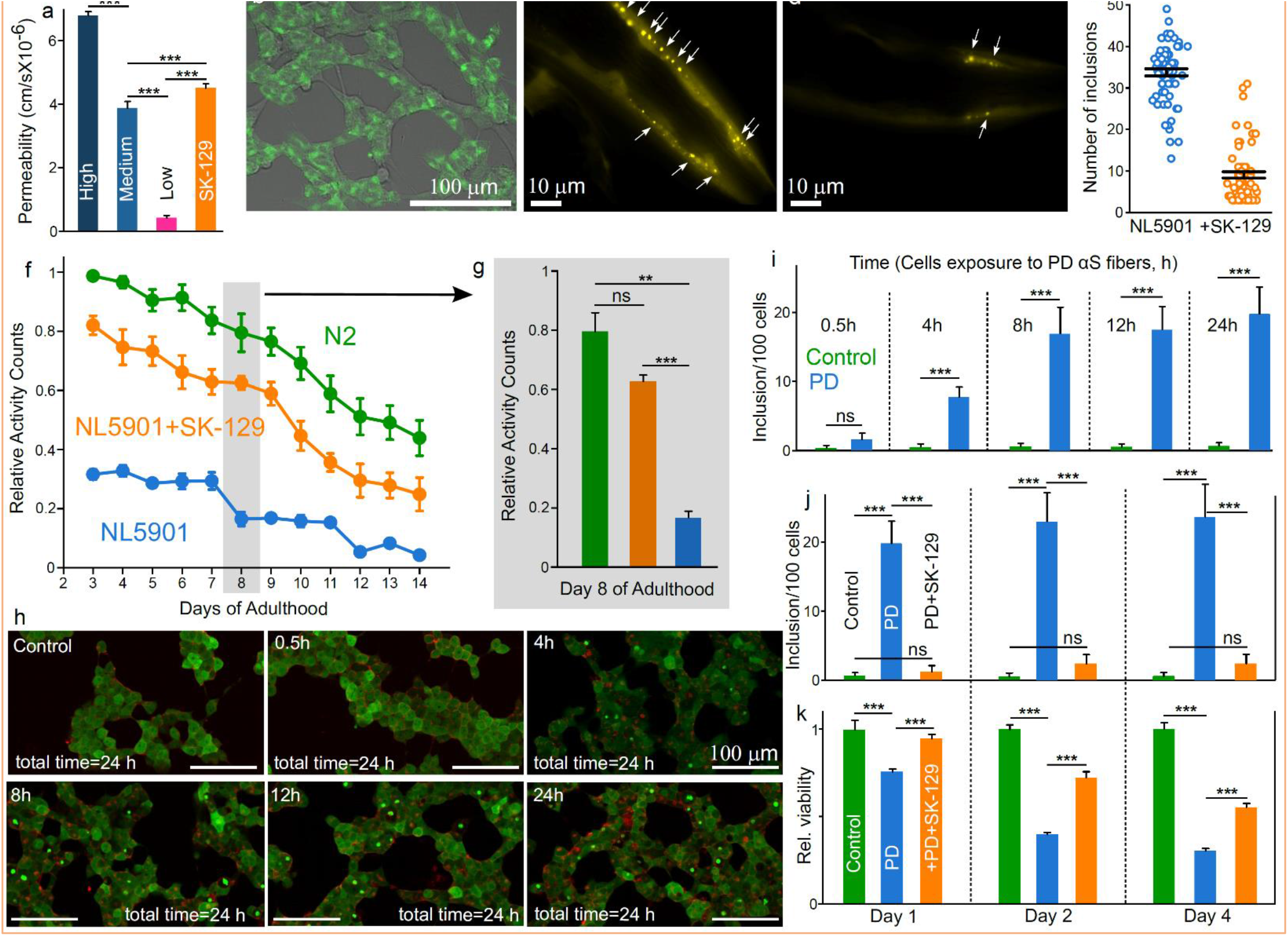
The intracellular inhibition of αS aggregation by SK-129 in PD models. Assessment of cell permeability of SK-129 using PAMPA (**a**) and confocal microscopy (**b**). Confocal image of SH-SY5Y cells after treatment with 100 nM SK-129F for 15 h. Confocal images (**c**,**d**) of αS-YFP inclusions (white arrows) in muscle cells of NL5901 (Days of adulthood=8 days) in the absence (**c**) and presence (**d**) of 15 µM SK-129. **e**, The number of inclusions for experiment ‘**c-d**’ for NL5901 in the absence and presence of SK-129. **f**, The locomotion for 14 days of adulthood of N2 and NL5901 and statistics (day 8) in the absence and presence of 15 µM SK-129. **h**, Confocal images and number of inclusions (**i**) of HEK cells incubated with PMCA sample from control and PD brain extract for indicated times. The HEK cells were washed after incubating them with the PMCA sample for various amounts of time. The number of inclusions (**j**) and relative viability (**k**) when HEK cells were incubated with the PMCA sample from PD brain extract for 8 h followed by washing, and further incubating cells up to 4 days. ^*^p<0.05, ^**^p<0.01, ***p<0.001, ****p<0.0001.

### Effect of SK-129 on the intracellular prion-like spread of αS seeds

SK-129 was potent in inhibiting *in vitro* prion-like spread of αS seeds. We developed a novel assay to test the antagonist activity of SK-129 in an intracellular prion-like spread model of αS using HEK cells. First, we assessed the total time required by αS seeds for the internalization into HEK cells. The HEK cells (expressing αS-A53T-YFP) were exposed to αS seeds (0.125 µM) extracted from PD brain for various time points (0.5, 4, 8, 12, and 24 h), washed the cells, and incubated for a total of 24 h. The formation of αS inclusions was noticeable within 4 h treatment of cells with αS seeds and inclusions were comparable (∼20 inclusions/100 cells) in the case of 8,12, and 24 h treatment of cells (Fig. 5h,i), which suggests that αS seeds were completely internalized in cells within 8 h. The antagonist activity of SK-129 against the intracellular prion-like spread was measured using the HEK cells treated with αS seeds for 8 h. SK-129 (10 µM) was added to HEK cells that were already treated with αS seeds (0.125 µM) for 8 h, followed by the incubation for an additional 16 h (total 24 h) (Fig. 5j). There was an abundance of inclusions in the absence of SK-129 after 24 h (∼19 inclusions/100 cells, Fig. 5i); however, a low number of inclusions (∼1-2 inclusions/100 cells) were observed in the presence of SK-129 (Fig. 5j). We also observed a gradual increase in inclusions and cytotoxicity in the presence of αS seeds for up to 4 days. However, in the presence of SK-129, a low number of inclusions was observed for up to 4 days (2-3 inclusions/100 cells, Fig. 5j), and the cell viability was improved significantly up to four days (Fig. 5k). Therefore, we conclude that SK-129 is a potent inhibitor of the intracellular *de novo* αS aggregation and the prion-like spread of αS seeds.

## DISCUSSION

αS aggregation is one of the causal agents in PD pathologies, making it an enticing therapeutic target. However, the atomic-level understanding of the sequences that initiate αS aggregation is limited; therefore, strategies that identify aggregation-prone αS sequences could have significant therapeutic implications in the treatment of PD. We used OQs as a multipronged approach to investigate αS aggregation on a molecular level and to identify novel targets that are essential for the initiation of αS aggregation.

The study led to the identification of SK-129 as a potent inhibitor of αS aggregation under both *in vitro* and *in vivo* PD models. The data suggest that SK-129 stabilizes αS in an amyloid incompetent helical conformation by specifically interacting with distinct αS sequences. Deletion of these sequences completely abolished the *de novo* and seed-catalyzed aggregation of αS. We postulate that the identified αS sequences are essential to initiate the aggregation and could be considered as novel therapeutic targets for the potent inhibition of αS aggregation. In groundbreaking findings, Eisenberg^13^ and Radford^12^ groups have identified αS sequences that initiate aggregation and they are in close proximity to the αS sequences identified from our study. More importantly, we have also validated these αS sequences by targeting them with foldamers, which led to the complete inhibition of αS aggregation and rescue of PD phenotypes in vivo and in vitro models. Our data suggest that the aggregation-prone αS sequences are potentially sampling helical conformation during αS aggregation; therefore, the design of helical mimetics complementing the chemical fingerprints of the helical conformation of αS sequences could lead to effective antagonism of αS aggregation and rescue of PD phenotypes.

We demonstrated that αS sequences that initiate *de novo* αS aggregation are important for the seed-catalyzed aggregation, which mimics the prion-like spread of αS. The prion-like spread requires the interaction of αS fibers with αS monomers to accelerate and propagate the aggregation. We surmise that SK-129 modulates αS monomer into fiber-incompetent conformation, which consequently prevents the prion-like spread of αS. We also developed a novel intracellular prion-like spread model of αS seeds (extracted from a PD brain). SK-129 was very effective in inhibiting the intracellular prion-like spread of αS seeds.

Our atomic-level study suggests that SK-129 regulates αS aggregation by binding to N-terminal sequences of αS, which are in close vicinity to the binding sites of molecular chaperones (Fig. 2, secB)^60^. Our NMR and CD data suggest that SK-129 regulates αS aggregation by shifting the equilibrium toward soluble and potentially functional αS, similar to molecular chaperones. The chaperones have been shown to interact with the N-terminal region of αS and shift the conformational equilibrium towards the functional αS to maintain the cellular homeostatic balance ^60^. The modulation of αS aggregation by affibody^61^ and molecular chaperones^60^ could be an attractive therapeutic intervention for PD; however, proteins/peptides are limited with poor cell permeability and poor enzymatic and conformational stability in biological milieus. SK-129 has demonstrated chaperone/affibody-like ability to manipulate αS aggregation as well as good pharmaceutical properties. More importantly, the side chains on SK-129 based scaffold can be conveniently manipulated without disturbing its overall conformation for further optimization of activity, which is often challenging with proteins/peptides.

To the best of our knowledge, this is the first report that simultaneously led to the identification and validation of the chemical fingerprints of key sequences, which initiate αS aggregation and the targeting of these sequences completely abolishes αS aggregation.

## METHODS

The methods and the supporting information sections include (1) protein expression (WT αS and mutants); (2) synthetic protocol and characterization of novel OQs, including SK-129F; (3) HSQC NMR’s between αS and SK-129 and LUVs; (4) the mutation study, including fluorescence polarization titrations between αS mutants and SK-129, ThT aggregation assays, TEMs, and seed-catalyzed aggregation of αS mutants; (5) HEK cells based seed catalyzed intracellular aggregation and toxicity assays; (6) extraction of αS seeds from the substantia nigra of the PD brain; (7) locomotion and confocal microscopy-based experiments to assess PD phenotypic readouts in *C elegans* PD model in the absence and presence of SK-129; (8) movie of the confocal images (3 D stack) of the NL5901 strain in the absence (Movie S1) and presence of SK-129 (Movie S2); and (8) all *in vitro* and *in vivo* studies presented in the manuscript.

## ACKNOLEDGMENTS

The authors like to thank the department of chemistry and biochemistry, The Knoebel Institute for Healthy Aging, and the University of Denver for the startup up funds. We sincerely thank The Fitch lab at NYU, Department of Biology, especially Prof. David Fitch and Dr. Karin Kiontke, for providing training to Sunil Kumar, which helped him in establishing *C elegans* based PD system in his lab. We also thank Prof. Lotta Granholm-Bentley, director of the Knoebel Institute for Healthy Aging, for the comments and proofreading of this manuscript. The author also thanks the PinS program (University of Denver) for awarding summer undergraduate fellowships to T.C.F., C.M.D., and M.M.B.

## ADDITIONAL INFORMATION

Correspondence and requests for materials should be addressed to S.K.

## AUTHOR CONTRIBUTIONS

S.K. designed and conceived the project with assistance from J.A. The synthesis of OQs and their derivatives were carried out by S.K. and J.A. The biophysical study was carried out by J.A. with some assistance from S.K. The NMR study was carried out by J.A. with some assistance from S.K. The design, expression, and purification of the αS mutants and the WT αS was carried out by J.A. The biophysical characterization of αS mutants was carried out by J.A. The SH-SY5Y cell toxicity assays were carried out by T.C.F. The HEK cell-based cytotoxicity and confocal microscopy imaging were carried out by C.D. with initial assistance from T.C.F. The extraction of the αS seeds from the post mortem PD brain was carried out by J.A. with the help from T.C.F. The *C elegans-*based *in vivo* experiments to monitor the locomotion were carried out by J.J. The confocal microscopy imaging experiments with *C elegans* and HEK cells were conducted by S.K. and C.D. The primary hippocampal neuron experiments were carried out by J.A. with the help of C.Z. from Y. Q.’s lab. The toxicity assays for primary hippocampal neurons were carried out by J.A. and T.C.F. The confocal microscopy imaging and immunochemistry experiments with the primary hippocampal neurons were carried out by J.A. with assistance from S.K. The flow cytometry experiments were conducted by S.K. and A.S., with assistance from S.H. The paper was written by S.K. with assistance from J.A., with editing from S.H.

## CITATIONS

1) Spillantini, M. G., Schmidt, M. L., Lee, V. M., Trojanowski, J. Q., Jakes, R. & Goedert, M. Alpha-synuclein in Lewy bodies. Nature 388, 839–840 (1997).

2) Dettmer, U., Selkoe, D. & Bartels, T. New insights into cellular α-synuclein homeostasis in health and disease. Curr. Opin. Neurobiol. 36, 15–22 (2016).

3) Chiti, F. & Dobson, C. M. Protein misfolding, functional amyloid, and human disease. Annu. Rev. Biochem. 75, 333–366 (2006).

4) Dawson, T. M. & Dawson, V. L. Molecular pathways of neurodegeneration in Parkinson’s disease. Science 302, 819–822 (2003).

5) Goedert, M. Alpha-synuclein and neurodegenerative diseases. Nat. Rev. Neurosci. 2, 492–501 (2001).

6) Buell, A. K., Galvagnion, C., Gaspar, R., Sparr, E., Vendruscolo, M., Knowles, T. P., Linse, S. & Dobson, C. M. Solution conditions determine the relative importance of nucleation and growth processes in α-synuclein aggregation. Proc. Natl. Acad. Sci. USA 111, 7671–7676 (2014).

7) Fink, A. L. The aggregation and fibrillation of alpha-synuclein. Acc. Chem. Res. 39, 628–634 (2006).

8) Burré, J., Sharma, M. & Südhof, T. C. Cell Biology and Pathophysiology of α-Synuclein. Cold Spring Harb. Perspect. Med. 8, a024091 (2018).

9) Meade, R. M., Fairlie, D. P. & Mason, J. M. Alpha-synuclein structure and Parkinson’s disease - lessons and emerging principles. Mol. Neurodegener. 14, 29 (2019).

10) Kingwell, K. Zeroing in on neurodegenerative α-synuclein. Nat. Rev. Drug Discov. 16, 371–373 (2017).

11) Pujols, J., Peña-Díaz, S., Pallarès, I. & Ventura, S. Chemical Chaperones as Novel Drugs for Parkinson’s Disease. Trends. Mol. Med. 26, 408–421 (2020).

12) Doherty, C., Ulamec, S. M., Maya-Martinez, R., Good, S. C., Makepeace, J., Khan, G. N., Van Oosten-Hawle, P., Radford, S. E. & Brockwell, D. J. A short motif in the N-terminal region of α-synuclein is critical for both aggregation and function. Nat. Str. Mol. Biol. 27, 249–259 (2020).

13) Rodriguez, J. A., Ivanova, M. I., Sawaya, M. R., Cascio, D., Reyes, F. E., Shi, D., Sangwan, S., Guenther, E. L., Johnson, L. M., Zhang, M., Jiang, L., Arbing, M. A., Nannenga, B. L., Hattne, J., Whitelegge, J., Brewster, A. S., Messerschmidt, M., Boutet, S., Sauter, N. K., Gonen, T. & Eisenberg, D. S. Structure of the toxic core of α-synuclein from invisible crystals. Nature 525, 486–490 (2015).

14) Crowther, R. A., Jakes, R., Spillantini, M. G. & Goedert, M. Synthetic filaments assembled from C-terminally truncated alpha-synuclein. FEBS lett. 436, 309–312 (1998).

15) Kessler, J. C., Rochet, J. C. & Lansbury, P. T. The N-terminal repeat domain of alpha-synuclein inhibits beta-sheet and amyloid fibril formation. Biochemistry 42, 672–678 (2003).

16) Izawa, Y., Tateno, H., Kameda, H., Hirakawa, K., Hato, K., Yagi, H., Hongo, K., Mizobata, T., & Kawata, Y. Role of C-terminal negative charges and tyrosine residues in fibril formation of α-synuclein. Brain Behav. 2, 595–605 (2012).

17) Guichard, G. & Huc, I. Synthetic foldamers. Chem Commun (Camb). 47, 21 (2011).

18) Chandramouli, N., Ferrand, Y., Lautrette, G., Kauffmann, B., Mackereth, C. D., Laguerre, M., Dubreuil, D. & Huc, I. Iterative design of a helically folded aromatic oligoamide sequence for the selective encapsulation of fructose. Nat. Chem. 4, 334–341 (2015).

19) Collie, G. W., Pulka-Ziach, K., Lombardo, C. M., Fremaux, J., Rosu, F., Decossas, M., Mauran, L., Lambert, O., Gabelica, V., Mackereth, C. D. & Guichard, G. Shaping quaternary assemblies of water-soluble non-peptide helical foldamers by sequence manipulation. Nat. Chem. 7, 871−878 (2015).

20) Goodman, C. M., Choi, S., Shandler, S. & DeGrado, W. F. Foldamers as versatile frameworks for the design and evolution of function. Nat. Chem. Biol. 3, 252−262 (2007).

21) Kumar, S., Henning-Knechtel, A., Chehade, I., Magzoub, M. & Hamilton, A. D. Foldamer-Mediated Structural Rearrangement Attenuates Aβ Oligomerization and Cytotoxicity. J. Am. Chem. Soc. 139, 17098−17108 (2017).

22) Kumar, S., Birol, M., Schlamadinger, D. E., Wojcik, S. P., Rhoades, E. & Miranker, A. D. Foldamer-mediated manipulation of a pre-amyloid toxin. Nat. Commun. 7, 11412 (2016).

23) Kumar, S. & Miranker, A. D. A foldamer approach to targeting membrane bound helical states of islet amyloid polypeptide. Chem. Commun. (Camb). 49, 4749–4751 (2013).

24) Kumar, S., Brown, M. A., Nath, A. & Miranker, A. D. Folded small molecule manipulation of islet amyloid polypeptide. Chem. Biol. 21, 775–781(2014).

25) Kumar, S., Birol, M. & Miranker, A. D. Foldamer scaffold suggest distinct structures are associated with alternative gain-of-function in a pre-amyloid toxin. Chem. Commun. (Camb). 52, 6391–6394 (2016).

26) Birol, M., Kumar, S., Rhoades, E. & Miranker, A. D. Conformational switching within dynamic oligomers underpins toxic gain-of-function by diabetes-associated amyloid. Nat. Commun. 9, 1312 (2018).

27) Levine, H. Thioflavine T interaction with synthetic Alzheimer’s disease β-amyloid peptides: detection of amyloid aggregation in solution. Protein Sci. 2, 404−410 (1993).

28) Bieschke, J., Russ, J., Friedrich, R. P., Ehrnhoefer, D. E. & Wobst, H., Neugebauer, K. & Wanker, E. E. EGCG remodels mature α-synuclein and amyloid-β fibrils and reduces cellular toxicity. Proc. Natl. Acad. Sci. USA. 107, 7710–7715 (2010).

29) Gurry, T., Ullman, O., Fisher, C. K., Perovic, I., Pochapsky, T. & Stultz, C. M. The Dynamic Structure of α-Synuclein Multimers. J. Am. Chem. Soc. 135, 3865–3872 (2015).

30) Lashuel, H. A., Overk, C. R., Oueslati, A. & Masliah, E. The many faces of α-synuclein: from structure and toxicity to therapeutic target. Nat. Rev. Neurosci. 14, 38–48 (2013).

31) Xicoy H., Wieringa, B. & Martens, G. J. The SH-SY5Y cell line in Parkinson’s disease research: a systematic review. Mol. Neurodegen. 12, 10 (2017).

32) Sanders, D. W., Kaufman, S. K., DeVos, S. L., Sharma, A. M., Mirbaha, H., Li, A., Barker, S. J., Foley, A. C., Thorpe, J. R., Serpell, L. C., Miller, T. M., Grinberg, L. T., Seeley, W. W. & Diamond, M. I. Distinct tau prion strains propagate in cells and mice and define different tauopathies. Neuron 82, 1271– 88 (2014).

33) Sangwan, S., Sahay, S., Murray, K. A., Morgan, S., Guenther, E. L., Jiang, L., Williams, C. K., Vinters, H. V., Goedert, M. & Eisenberg, D. S. Inhibition of synucleinopathic seeding by rationally designed inhibitors. eLife 9, e46775 (2020).

34) Probst, C. Characterization of Protein Aggregates, Silicone Oil Droplets, and Protein-Silicone Interactions Using Imaging Flow Cytometry. J. Pharm. Sci. 109, 364–74 (2020).

35) Runwal, G. & Edwards, R. H. The membrane interactions of synuclein: physiology and pathology. Annu. Rev. Pathol. Mech. Dis. 16, 465–85 (2021).

36) Fusco, G., Simone, A. D., Gopinath, T., Vostrikov, V., Vendruscolo, M., Dobson, C. M. & Veglia, G. Direct observation of the three regions in α-synuclein that determine its membrane-bound behaviour. Nat. Commun. 5, 3827 (2014).

37) Galvagnion, C., Buell, A. K., Meisl, G., Michaels, T. C. T., Vendruscolo, M., Knowles, T. P. J. & Dobson, C. M. Lipid vesicles trigger α-synuclein aggregation by stimulating primary nucleation. Nat. Chem. Biol. 3, 229–34 (2015).

38) Galvagnion, C., Brown, J. W. P., Oluberai, M. M., Flagmeier, P., Vendruscolo, M., Buell, A. K., Sparr, E. & Dobson, C. M. Chemical properties of lipids strongly affect the kinetics of the membrane-induced aggregation of α-synuclein. Proc. Natl. Acad. Sci. USA 26, 7065–70 (2016).

39) Ulmer, T. S, Bax A., Cole, N. B. & Nussbaum, R. L., Structure and Dynamics of Micelle-bound Human α-Synuclein. J. Biol. Chem. 280, 9595–603 (2005).

40) Fusco, G., Chen, S. W., Williamson P. T. F., Cascella R., Perni M., Jarvis J.A., Cecchi C., Vendruscolo M., Chiti F., Cremades N., Ying L., Dobson C.M. & Simone A. D. Structural basis of membrane disruption and cellular toxicity by a-synuclein oligomers. Science 358, 1440–3 (2017).

41) Goedert, M., Falcon B., Clavaguera, F. & Tolnay, M. Prion-like Mechanisms in the Pathogenesis of Tauopathies and Synucleinopathies. Curr Neurol and Neurosci Rep. 14, 495 (2014).

42) Goedert, M. Alzheimer’s and Parkinson’s diseases: The prion concept in relation to assembled Aβ, tau, and α-synuclein. Science 349, 1255555 (2015).

43) Braak, H., Tredici, K. D., Rüb, U., Vos, R. A. D., Steur, E. N. J. & Braak, E. Staging of brain pathology related to sporadic Parkinson’s disease. Neurobiol. Aging 24, 197–211 (2003).

44) Braak, H. & & Tredici, K.D. Neuroanatomy and Pathology of Sporadic Parkinson’s Disease. Adv. Anat. Embryol. Cell Biol. 201, 1–119 (2009).

45) Luk, K. C., Song, C., O’Brien, P., Stieber, A., Branch, J. R., Brunden, K. R., Trojanowski, J. Q. & Lee, V. M. Y. Exogenous α-synuclein fibrils seed the formation of Lewy body-like intracellular inclusions in cultured cells. Proc. Natl. Acad. Sci. USA 106, 20051–6 (2009).

46) Masuda-Suzukake, M., Nonaka, T., Hosokawa, M., Oikawa, T., Arai, T., Akiyama, H., Mann, D. M. A. & Hasegawa, M. Prion-like spreading of pathological α-synuclein in brain. Brain 136, 1128–1138 (2013).

47) Desplats, P., Lee, H. J., Bae, E. J., Patrick, C., Rockenstein, E., Crews, L., Spencer, B., Masliah, E. & Lee, S. J. Inclusion formation and neuronal cell death through neuron-to-neuron transmission of α-synuclein. Proc. Natl. Acad. Sci. USA 106, 13010–13015 (2009).

48) Mahul-Mellier, A., Burtscher, J., Maharjan, N., Weerens, L., Croisier, M., Kuttler, F., Leleu, M., Knott, G. W. & Lashuel H. A. The process of Lewy body formation, rather than simply α-synuclein fibrillization, is one of the major drivers of neurodegeneration. Proc. Natl. Acad. Sci. USA 117, 4971–4982 (2020).

49) Pujols, J., Peña-Díaz, S., Lázaro, D. F., Peccati, F., Pinheiro, F., González, D., Carija, A., Navarro, S., Conde-Giménez, M., García, J., Guardiola, S., Giralt, E., Salvatella, X., Sancho, J., Sodupe, M., Outeiro, T.F., Dalfó, E. & Ventura, S. Small molecules inhibits a-synuclein aggregation, disrupts amyloid fibrils, and prevents degeneration of dopaminergic neurons. Proc. Natl. Acad. Sci. USA 115, 10481–10486 (2018).

50) Barria, M. A., Gonzalez-Romero, D. & Soto, C. Cyclic amplification of prion protein misfolding. Methods Mol. Biol. 849, 199–212 (2012).

51) Shahnawaz, M., Mukherjee, A., Pritzkow, S., Mendez, N., Rabadia, P., Liu, X., Hu, B., Schmeichel, A., Singer, W., Wu, G., Tsai, A.L., Shirani, H., Nilsson, K.P.R., Low, P.A. & Soto, C. Discriminating α-synuclein strains in Parkinson’s disease and multiple system atrophy. Nature 578, 273–277 (2020).

52) Strohäker, T., Jung, B. C., Liou, S. H., Fernandez, C. O., Riedel, D., Becker, S., Halliday, G. M., Bennati, M., Kim, W. S., Lee, S. J. & Zweckstetter, M. Structural heterogeneity of α-synuclein fibrils amplified from patient brain extracts. Nat. Commun. 10, 5535 (2019).

53) Bousset, L., Pieri, L., Ruiz-Arlandis, G., Gath, J., Jensen, P. H., Habenstein, B., Madiona, K., Olieric, V., Böckmann, A., Meier, B. H. & Melki, R. Structural and functional characterization of two alpha-synuclein strains. Nat. Commun. 4, 2575 (2013).

54) Jameson, L.P., Smith, N.W. & Dzyuba, S.V. Dye-binding assays for evaluation of the effects of small molecule inhibitors on amyloid (aβ) self-assembly. ACS. Chem. Neurosci. 3, 807–819 (2012).

55) Tanik, S. A., Schultheiss, C. E., Volpicelli-Daley, L. A., Brunden, K. R. & Lee, V. M. Lewy body-like α-synuclein aggregates resist degradation and impair macroautophagy. J. Biol. Chem. 288, 15194–15210 (2013).

56) Volpicelli-Daley, L. A., Luk, K. C., Patel, T. P., Tanik, S. A., Riddle, D. M., Stieber, A., Meaney, D. F., Trojanowski, J. Q. & Lee V. M. Exogenous α-synuclein fibrils induce Lewy body pathology leading to synaptic dysfunction and neuron death. Neuron 72, 57–71 (2011).

57) Van Ham, T. J., Thijssen, K. L., Breitling, R., Hofstra, R. M., Plasterk, R. H. & Nollen, E. A. C. elegans model identifies genetic modifiers of alpha-synuclein inclusion formation during aging. PLoS Genet. 4, 1000027 (2008).

58) Currey, H. N., Malinkevich, A., Melquist, P. & Liachko, N. F. ARENA-based activity profiling of tau and TDP-43 transgenic C. elegans. MicroPubl. Biol. (2020).

59) Garcia-Moreno, J.C., Porta de la Riva, M., Martínez-Lara, E., Siles, E. & Cañuelo, A. Tyrosol, a simple phenol from EVOO, targets multiple pathogenic mechanisms of neurodegeneration in a C. elegans model of Parkinson’s disease. Neurobiol. Aging 82, 60–68 (2019).

60) Burmann, B. M., Gerez, J.A., Matecko-Burmann, I., Campioni, S., Kumari, P., Ghosh, D., Mazur, A., Aspholm, E.E., Sulskis, D., Wawrzyniuk, M., Bock, T., Schmidt, A., Rudiger, S.G.D., Riek, R. & Hiller, S. Regulation of α-synuclein by chaperones in mammalian cells. Nature 577, 127–132 (2020).

61) Agerschou, E. D., Flagmeier, P., Galvagnion, C., Komnig, D., Heid, L., Prasa, V., Shaykhalishahi, H. & Willbold, D. An engineered monomer binding-protein for α-synuclein efficiently inhibits the proliferation of amyloid fibrils. eLife 9, e46112 (2019).

