## Supplementary information for "Foldamers Reveal and Validate Novel Therapeutic Targets Associated with Toxic α-Synuclein Self-Assembly"

*I would like to dedicate this paper to Professor Andrew D. Hamilton, President of New York University, who has contributed immensely to the field of chemical biology, a great mentor, and a very humble human being.

Synthesis of compounds 1, 2, 3, 4, 6a, and 7 have been reported somewhere else[1,2].

**Standard Protocol for Reduction of Nitro-oligoquinolines**

To a solution of nitroquinoline (0.1-0.5 mmol) in ethylacetate/dichloromethane (DCM) (1:1, v,v, 10 mL), Pd/C (15% wt.) was added and the reaction started with constant stirring at room temperature in the atmosphere of $H_2$(g). The progress of the reaction was followed by TLC (Thin layer chromatography). The reaction was stopped at the disappearance of the starting material, which was around 15-20 h. The reaction mixture was filtered and dried, which result in a yellow solid with quantitative yield. The product was used in the next step without further characterization.

**Standard Protocol for Amide Coupling**

To a solution of nitroquinoline carboxylic acid (1.2 mmol) in DCM (10 mL, anhydrous), triethylamine (4.8 mmol, 4 mole equivalent, anhydrous) and 2-chloromethyl-1-methyl pyridinium iodide (1.5 mmol) were added and the reaction was refluxed for 20 min. at 50°C under an inert atmosphere of argon (g). To this solution, amino-oligoquinoline (1.0 mmol) in 3 mL DCM (anhydrous) was added and the reaction started with constant stirring at 50°C under an inert atmosphere. The reaction mixture was stirred for 6 h after which the volatiles were removed on the rotovap. Flash chromatography (0 to 45% ethyl acetate in hexane, v/v) yielded the desired product as a yellow to brown solid.

**Standard Protocol for Deprotection of Oligoquinolines**

To a solution of oligoquinoline (0.04 mmol), a cocktail solution (2 mL, DCM: trifluoroacetic acid (TFA): triethylsilane, 70:20:10, v/v) was added and the solution was stirred at room temperature for 4 h. The reaction mixture was dried and washed with cold diethyl ether (4 × 3mL), which results in a yellow to brown solid. The compound were redissolved in DMSO and purified using HPLC with buffer A (95% water, 5% acetonitrile, 0.1% TFA) and buffer B (95% acetonitrile, 5% water, 0.1% TFA). The gradient for buffer A to B was used from 100% to 0% for a total of 20 min. at a rate of 3 mL/min. on a reverse-phase C-18 semiprep column (Hypersil gold, 150 mm × 10 mm).

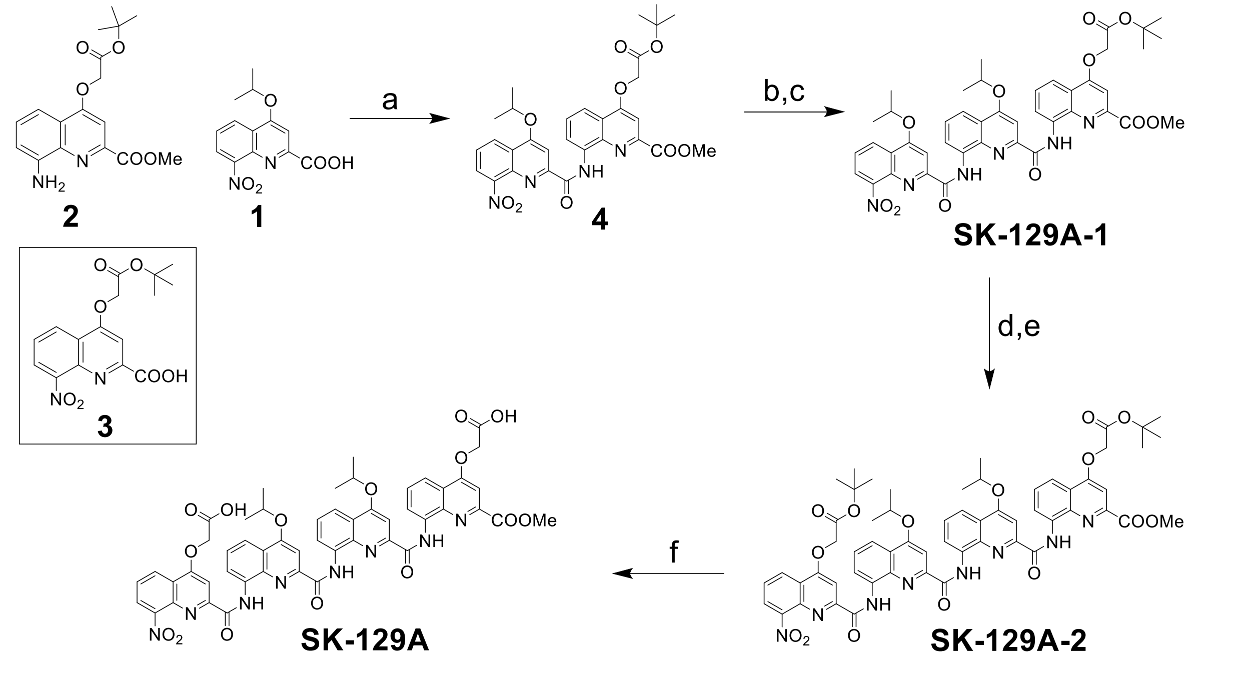


**Scheme 1**. The conditions for the synthesis of SK-129A. **a**, 2-chloro-1-methylpyridinium iodide, DCM (Anhydrous), trimethylamine (Anhydrous), 12 h, 50 °C. **b**, $H_2$(g), Pd/C, ethylacetate, 12 h, r.t. **c**, **1** and 6, 2-chloro-1-methylpyridinium iodide, DCM, trimethylamine, 12 h, 50 °C. **d**, $H_2$(g), Pd/C, ethylacetate, 12 h, r.t. **e**, **3** and 6, 2-chloro-1-methylpyridinium iodide, DCM (Anhydrous), trimethylamine, 12 h, 50 °C. **f**, Triflouroacetic acid, DCM, triethylsilane, 4 h, r.t.

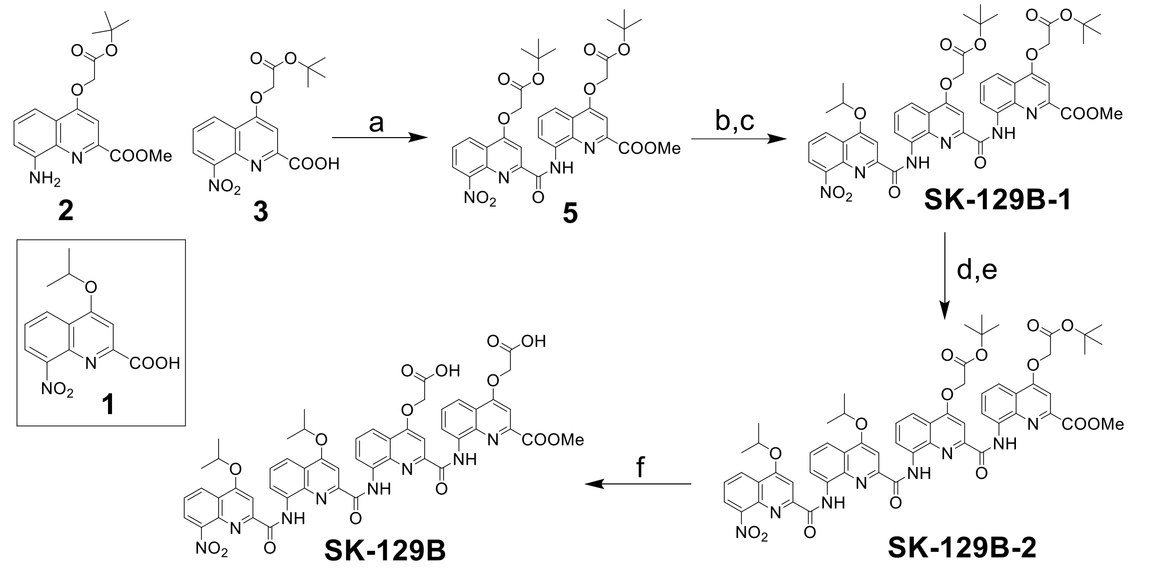


**Scheme 2**. The conditions for the synthesis of SK-129B. **a**, 2-chloro-1-methylpyridinium iodide, DCM (Anhydrous), trimethylamine (Anhydrous), 12 h, 50 °C. **b**, $H_2$(g), Pd/C, ethylacetate, 12 h, r.t. **c**, **1** and 6, 2-chloro-1-methylpyridinium iodide, DCM, trimethylamine, 12 h, 50 °C. **d**, $H_2$(g), Pd/C, ethylacetate, 12 h, r.t. **e**, **1** and 6, 2-chloro-1-methylpyridinium iodide, DCM (Anhydrous), trimethylamine, 12 h, 50 °C. **f**, Triflouroacetic acid, DCM, triethylsilane, 4 h, r.t.

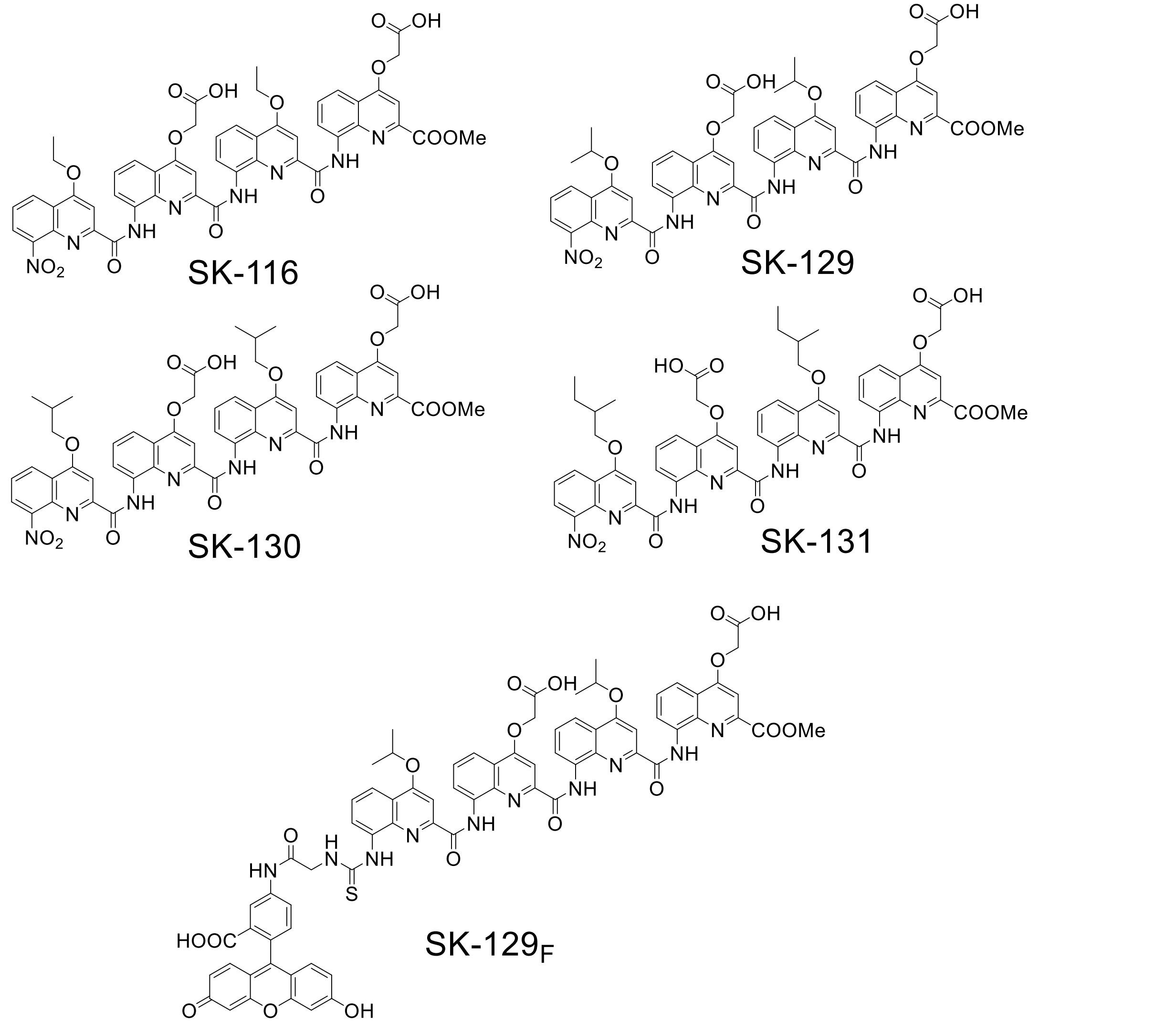


**Scheme 3**. Structures of the OQs used in the study.

**Synthesis of SK-129A-1**

$^{1}$H NMR (500 MHz, $CDCl_3$) δ 12.31 – 12.27 (s, 1H), 12.24 – 12.20 (s, 1H), 9.09 – 9.02 (ddd, *J* = 7.7, 4.7, 1.3 Hz, 2H), 8.48 – 8.42 (dd, *J* = 8.4, 1.5 Hz, 1H), 8.09 – 8.05 (dd, *J* = 8.4, 1.3 Hz, 1H), 8.05 – 8.01 (dd, *J* = 8.4, 1.3 Hz, 1H), 7.95 – 7.92 (s, 1H), 7.89 – 7.86 (s, 1H), 7.82 – 7.75 (t, *J* = 8.0 Hz, 1H), 7.71 – 7.64 (t, *J* = 8.1 Hz, 1H), 7.63 – 7.55 (dd, *J* = 7.5, 1.5 Hz, 1H), 7.44 – 7.37 (t, *J* = 7.9 Hz, 1H), 6.73 – 6.69 (s, 1H), 5.19 – 5.06 (dhept, *J* = 12.1, 6.1 Hz, 2H), 4.66 – 4.62 (s, 2H), 3.50 – 3.46 (s, 3H), 1.67 – 1.63 (d, *J* = 6.1 Hz, 6H), 1.60 – 1.56 (d, *J* = 6.1 Hz, 6H), 1.56 – 1.54 (s, 9H). MS (MALDI-TOF) calcd for $C_{43}H_{43}N_6O_{11}^{+}$ (M+H+): 819.30, obsd: 819.61.

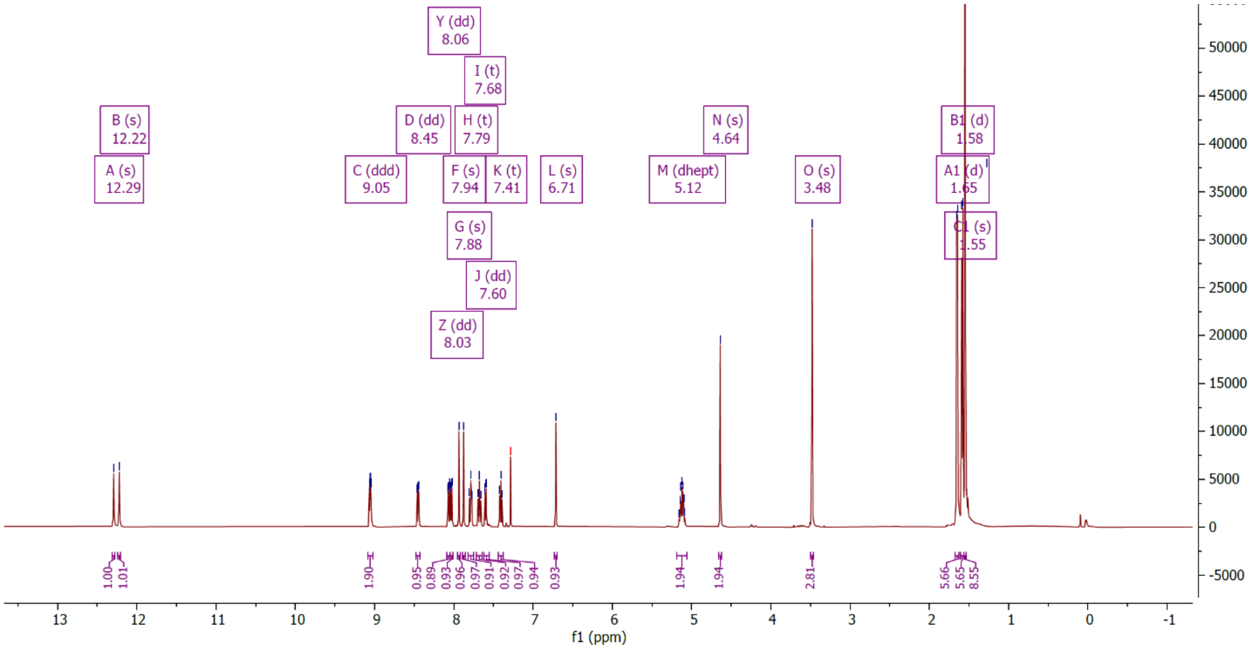

**Synthesis of SK-129A-2**

$^{1}$H NMR (500 MHz, $CDCl_3$) δ 12.41 – 12.37 (s, 1H), 11.95 – 11.91 (s, 1H), 11.73 – 11.69 (s, 1H), 9.18 – 9.12 (dd, *J* = 7.6, 1.2 Hz, 1H), 8.70 – 8.65 (dd, *J* = 8.3, 1.4 Hz, 1H), 8.49 – 8.43 (dd, *J* = 7.6, 1.2 Hz, 1H), 8.17 – 8.13 (dd, *J* = 7.6, 1.3 Hz, 1H), 8.13 – 8.10 (dd, *J* = 8.4, 1.2 Hz, 1H), 8.03 – 7.97 (dd, *J* = 8.4, 1.3 Hz, 1H), 7.95 – 7.89 (dd, *J* = 8.3, 1.3 Hz, 1H), 7.89 – 7.86 (s, 1H), 7.79 – 7.72 (t, *J* = 8.0, 8.0 Hz, 1H), 7.71 – 7.67 (t, *J* = 8.0, 8.0 Hz, 1H), 7.67 – 7.63 (dd, *J* = 7.5, 1.4 Hz, 1H), 7.47 – 7.40 (t, *J* = 7.9, 7.9 Hz, 1H), 7.36 – 7.29 (t, *J* = 8.0, 8.0 Hz, 1H), 6.91 – 6.87 (s, 1H), 6.65 – 6.62 (s, 1H), 5.18 – 5.08 (h, *J* = 6.1, 6.1, 6.1, 6.1, 6.1 Hz, 1H), 5.07 – 5.03 (s, 2H), 4.79 – 4.70 (h, *J* = 6.0, 6.0, 6.0, 6.0, 6.0 Hz, 1H), 4.68 – 4.64 (s, 2H), 3.50 – 3.46 (s, 3H), 1.70 – 1.65 (s, 6H), 1.65 – 1.62 (s, 9H), 1.57 – 1.53 (s, 9H), 1.53 – 1.46 (m, 6H). MS (MALDI-TOF) calcd for $C_{59}H_{59}N_8O_{15}^{+}$(M+H$^{+}$): 1119.41, obsd: 1120.04.

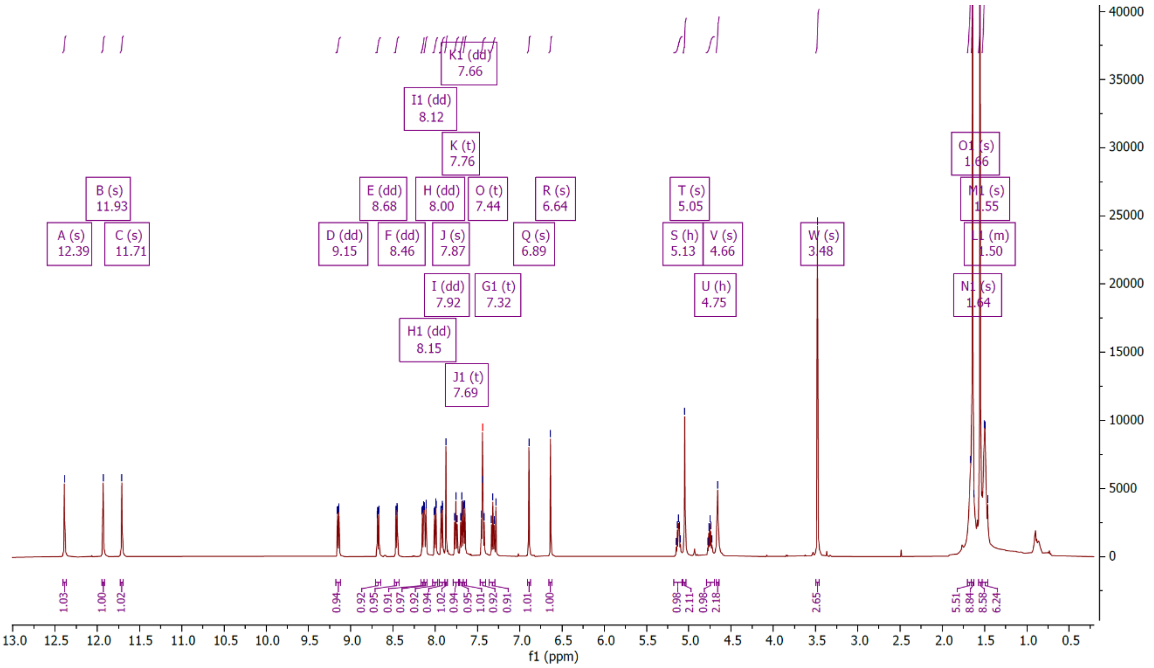

**Synthesis of SK-129A**

$^{1}$H NMR (500 MHz, DMSO) δ 12.20 – 12.16 (s, 1H), 11.73 – 11.70 (s, 1H), 11.47 – 11.44 (s, 1H), 9.06 – 9.00 (d, *J* = 7.6 Hz, 1H), 8.62 – 8.57 (d, *J* = 8.3 Hz, 1H), 8.41 – 8.36 (d, *J* = 7.6 Hz, 1H), 8.04 – 7.99 (d, *J* = 7.7 Hz, 1H), 7.98 – 7.91 (dd, *J* = 11.6, 8.4 Hz, 2H), 7.87 – 7.82 (d, *J* = 9.9 Hz, 2H), 7.82 – 7.74 (dq, *J* = 8.4, 4.4 Hz, 3H), 7.69 – 7.61 (t, *J* = 7.9 Hz, 1H), 7.48 – 7.40 (t, *J* = 8.1 Hz, 1H), 7.32 – 7.28 (s, 1H), 6.91 – 6.87 (s, 1H), 6.61 – 6.58 (s, 1H), 5.41 – 5.29 (m, 2H), 5.30 – 5.19 (h, *J* = 6.2 Hz, 1H), 5.04 – 4.92 (d, *J* = 16.5 Hz, 1H), 4.93 – 4.78 (dp, *J* = 12.2, 6.2 Hz, 2H), 3.39 – 3.35 (s, 3H), 1.72 – 1.58 (s, 3H), 1.58 – 1.51 (s, 4H), 1.51 – 1.35 (s, 6H). HRMS- ESI (m/z): calculated for $C_{51}H_{43}N_8O_{15}^{+}$[(M+H)$^{+}$]: 1007.2848, found 1007.2839. Anal. Calcd for $C_{51}H_{42}N_8O_{15}$: C, 60.83; H, 4.20; N, 11.13; O, 23.83. Found: C, 60.53; H, 4.34; N, 11.01.

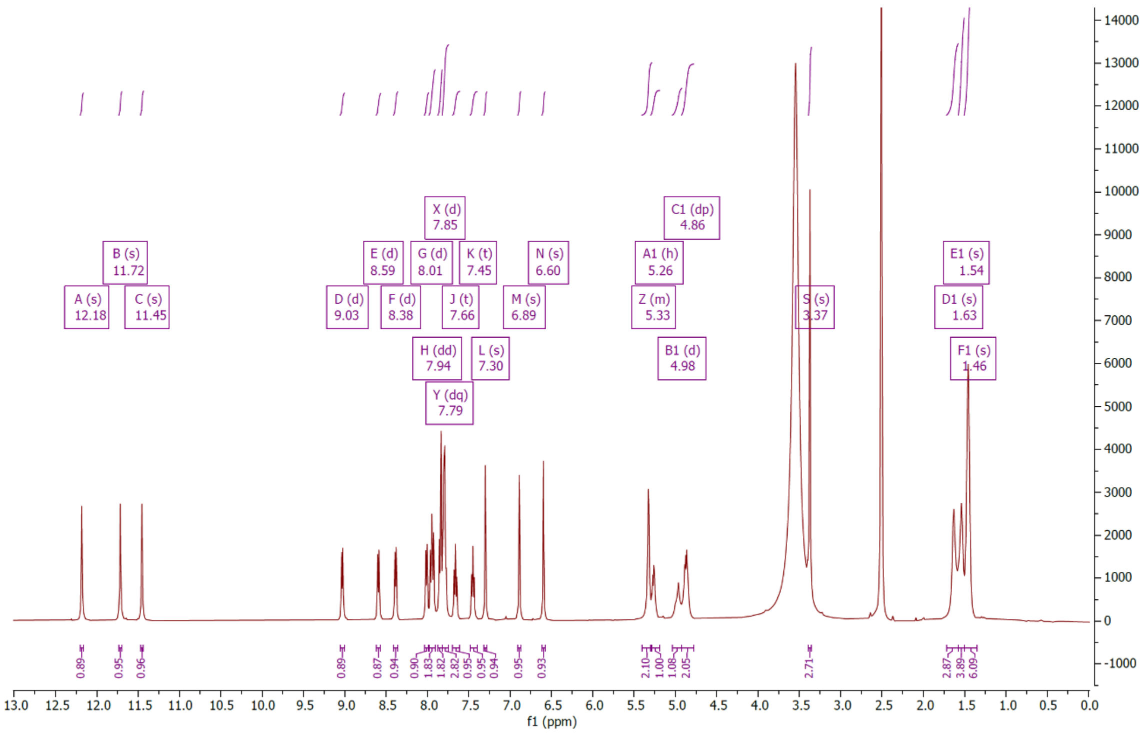

**Synthesis of SK-129B-1**

$^{1}H$ NMR (500 MHz, $CDCl_3$) δ 12.31 – 12.24 (s, 1H), 12.22 – 12.14 (s, 1H), 9.12 – 9.06 (d, *J* = 7.7 Hz, 1H), 9.06 – 9.00 (d, *J* = 7.6 Hz, 1H), 8.51 – 8.44 (d, *J* = 8.6 Hz, 1H), 8.20 – 8.13 (d, *J* = 8.4 Hz, 1H), 8.07 – 8.01 (d, *J* = 8.3 Hz, 1H), 7.96 – 7.93 (s, 1H), 7.81 – 7.76 (d, *J* = 11.0 Hz, 2H), 7.76 – 7.71 (t, *J* = 8.0 Hz, 1H), 7.65 – 7.57 (d, *J* = 7.4 Hz, 1H), 7.45 – 7.38 (t, *J* = 7.9 Hz, 1H), 6.73 – 6.69 (s, 1H), 5.18 – 5.09 (d, *J* = 6.6 Hz, 1H), 4.99 – 4.94 (s, 2H), 4.71 – 4.58 (s, 2H), 3.49 – 3.46 (s, 3H), 1.68 – 1.63 (d, *J* = 6.1 Hz, 6H), 1.59 – 1.55 (s, 9H), 1.33 – 1.26 (s, 9H). MS (MALDI-TOF) calcd for $C_{46}H_{47}N_6O_{13}^{+}$(M+H$^+$), 891.32, obsd: 891.71.

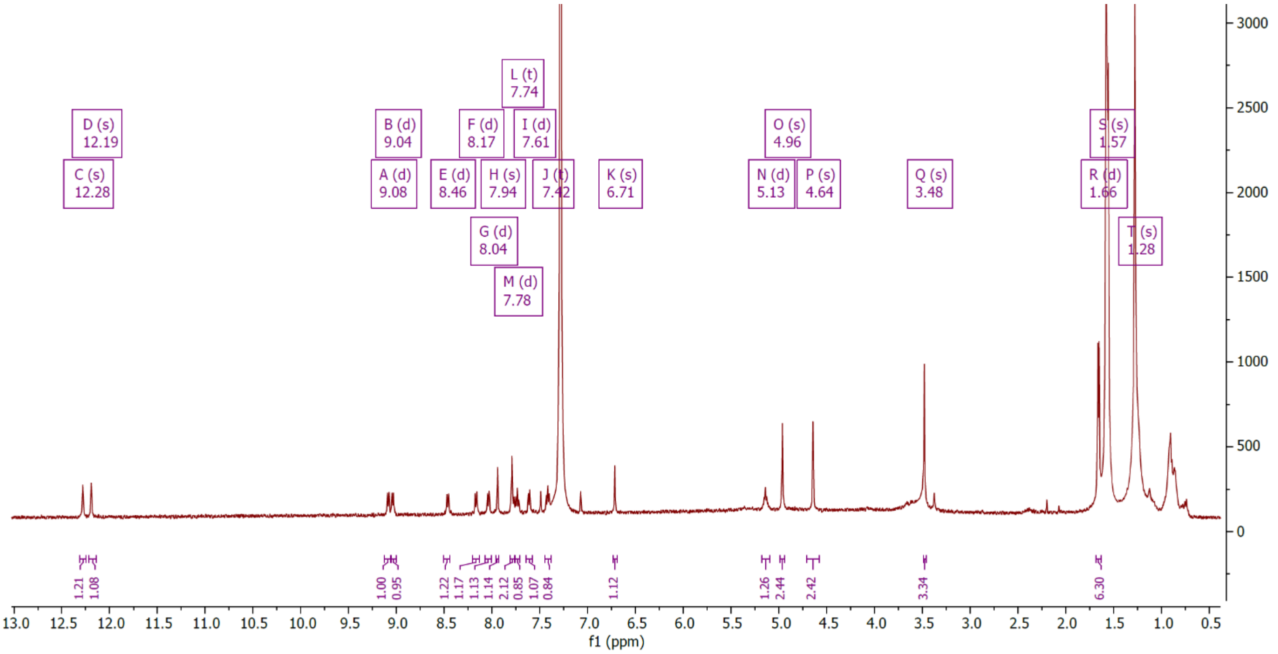

**Synthesis of SK-129B-2**

$^{1}H$ NMR (500 MHz, $CDCl_3$) δ 12.37 – 12.33 (s, 1H), 11.93 – 11.89 (s, 1H), 11.75 – 11.71 (s, 1H), 9.24 – 9.18 (dd, *J* = 7.7, 1.2 Hz, 1H), 8.59 – 8.48 (ddd, *J* = 24.5, 8.4, 1.5 Hz, 1H), 8.41 – 8.36 (dd, *J* = 7.6, 1.3 Hz, 1H), 8.23 – 8.18 (dd, *J* = 7.7, 1.3 Hz, 1H), 8.17 – 8.09 (dt, *J* = 8.4, 1.6 Hz, 2H), 8.02 – 7.88 (m, 2H), 7.85 – 7.79 (t, *J* = 8.0 Hz, 1H), 7.71 – 7.55 (m, 2H), 7.50 – 7.41 (m, 2H), 7.38 – 7.31 (t, *J* = 8.0 Hz, 1H), 6.88 – 6.84 (s, 1H), 6.67 – 6.63 (s, 1H), 5.18 – 5.06 (m, 2H), 4.74 – 4.70 (s, 2H), 4.69 – 4.65 (s, 2H), 3.48 – 3.45 (s, 3H), 1.70 – 1.64 (m, 7H), 1.59 – 1.56 (s, 10H), 1.56 – 1.55 (s, 8H), 1.54 – 1.50 (d, *J* = 6.0 Hz, 6H). MS (MALDI-TOF) calcd for $C_{59}H_{59}N_8O_{15}^{+}$(M+H$^{+}$), 1119.41, obsd: 1119.88.

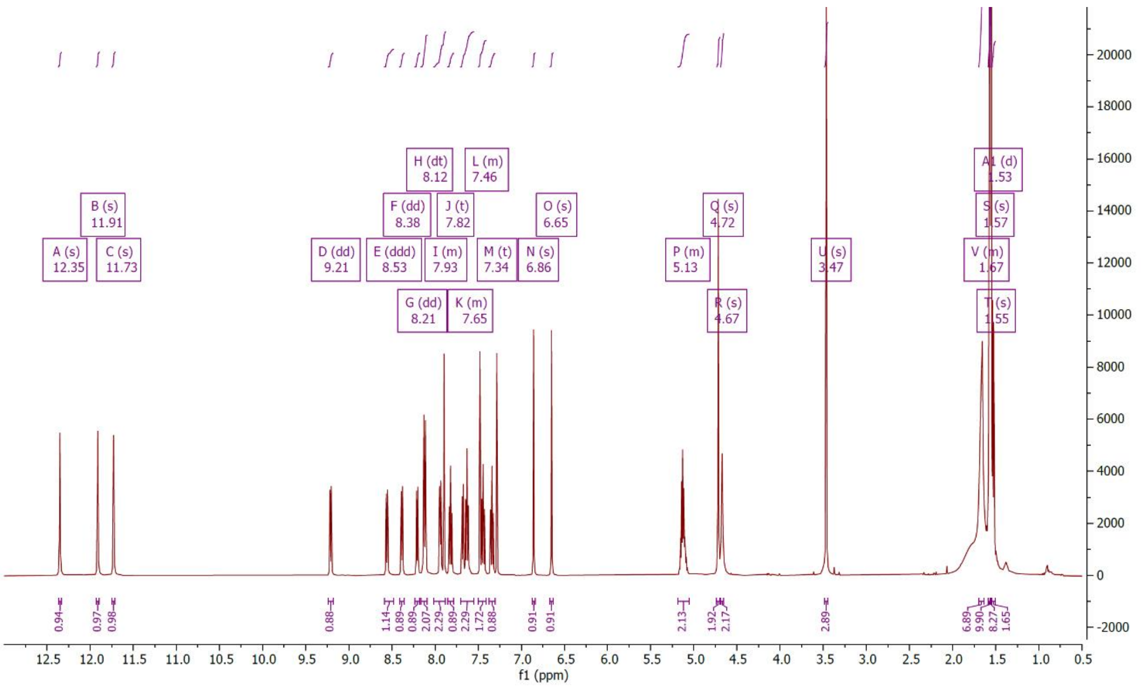

**Synthesis of SK-129B**

$^{1}$H NMR (500 MHz, DMSO) δ 12.13 – 12.09 (s, 1H), 11.72 – 11.69 (s, 1H), 11.51 – 11.47 (s, 1H), 9.11 – 9.06 (d, *J* = 7.6 Hz, 1H), 8.54 – 8.49 (d, *J* = 8.3 Hz, 1H), 8.36 – 8.31 (d, *J* = 7.6 Hz, 1H), 8.08 – 8.03 (d, *J* = 7.7 Hz, 1H), 8.03 – 7.98 (d, *J* = 8.4 Hz, 1H), 7.98 – 7.93 (d, *J* = 8.4 Hz, 1H), 7.89 – 7.83 (d, *J* = 8.1 Hz, 3H), 7.77 – 7.71 (t, *J* = 8.0 Hz, 1H), 7.71 – 7.67 (d, *J* = 7.4 Hz, 1H), 7.66 – 7.60 (t, *J* = 7.9 Hz, 1H), 7.50 – 7.43 (t, *J* = 8.0 Hz, 1H), 7.34 – 7.31 (s, 1H), 6.85 – 6.82 (s, 1H), 6.62 – 6.58 (s, 1H), 5.32 – 5.23 (pent, *J* = 6.1 Hz, 1H), 5.24 – 5.15 (pent, *J* = 6.0 Hz, 1H), 5.09 – 4.98 (s, 2H), 4.98 – 4.79 (d, *J* = 34.0 Hz, 2H), 3.45 – 3.30 (s, 3H), 1.92 – 1.72 (s, 3H), 1.69 – 1.59 (s, 4H), 1.59 – 1.47 (d, *J* = 29.8 Hz, 6H). HRMS-ESI (m/z): calculated for $C_{51}H_{43}N_8O_{15}^{+}$[$(M+H)^{+}$]: 1007.2848, found 1007.2842. Anal. Calcd for $C_{51}H_{42}N_8O_{15}$: C, 60.83; H, 4.20; N, 11.13; O, 23.83. Found: C, 60.59; H, 4.31; N, 11.08.

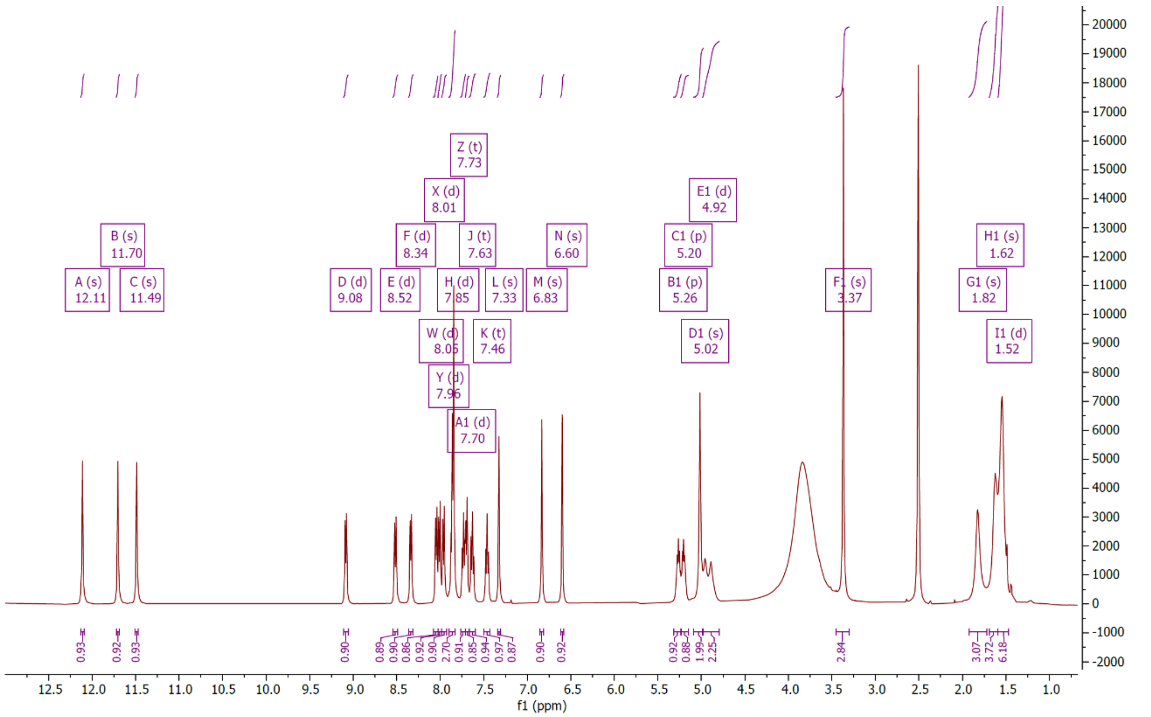

**Synthesis of SK-129-SCN**

To a solution of SK-129-$NH_2$ (30 mg, 0.027 mmol) in dichloromethane (10 mL), 1,1’-Thiocarbonyldi-2(1H)-pyridone (19.30 mg, 0.060 mmol, 3 equivalent) was added into the flask and the reaction solution was stirred for 12 h at room temperature in an inert atmosphere of argon. The progress of the reaction was monitored by TLC. Flash chromatography (0 to 40% ethylacetate in hexane) yielded the desired product as yellow solid (21 mg, 89%).

$^1$H NMR (500 MHz, $CDCl_3$) δ 12.63 – 12.60 (s, 1H), 12.01 – 11.95 (d, *J* = 11.6 Hz, 2H), 9.16 – 9.11 (d, *J* = 7.5 Hz, 1H), 8.51 – 8.46 (d, *J* = 7.5 Hz, 1H), 8.18 – 8.13 (d, *J* = 8.0 Hz, 2H), 8.12 – 8.07 (d, *J* = 8.4 Hz, 1H), 8.03 – 7.98 (d, *J* = 8.4 Hz, 2H), 7.79 – 7.73 (t, *J* = 7.9 Hz, 1H), 7.75 – 7.71 (s, 1H), 7.70 – 7.63 (t, *J* = 8.0 Hz, 1H), 7.44 – 7.41 (s, 1H), 7.40 – 7.34 (d, *J* = 8.0 Hz, 1H), 7.22 – 7.15 (t, *J* = 7.9 Hz, 1H), 6.85 – 6.82 (s, 1H), 6.66 – 6.63 (s, 1H), 6.63 – 6.58 (d, *J* = 7.4 Hz, 1H), 5.15 – 5.06 (pent, *J* = 6.1 Hz, 1H), 4.99 – 4.95 (m, 2H), 4.78 – 4.71 (t, *J* = 6.2 Hz, 1H), 4.70 – 4.55 (m, 2H), 3.56 – 3.52 (s, 3H), 1.92 – 1.87 (d, *J* = 6.1 Hz, 3H), 1.66 – 1.61 (s, 9H), 1.60 – 1.58 (m, 17H), 1.57 – 1.54 (s, 9H), 1.46 – 1.42 (d, *J* = 6.1 Hz, 3H). MS (MALDI-TOF) calcd for $C_{60}H_{59}N_8O_{13}S^+$(M+$H^+$), 1131.39, obsd: 1131.91.

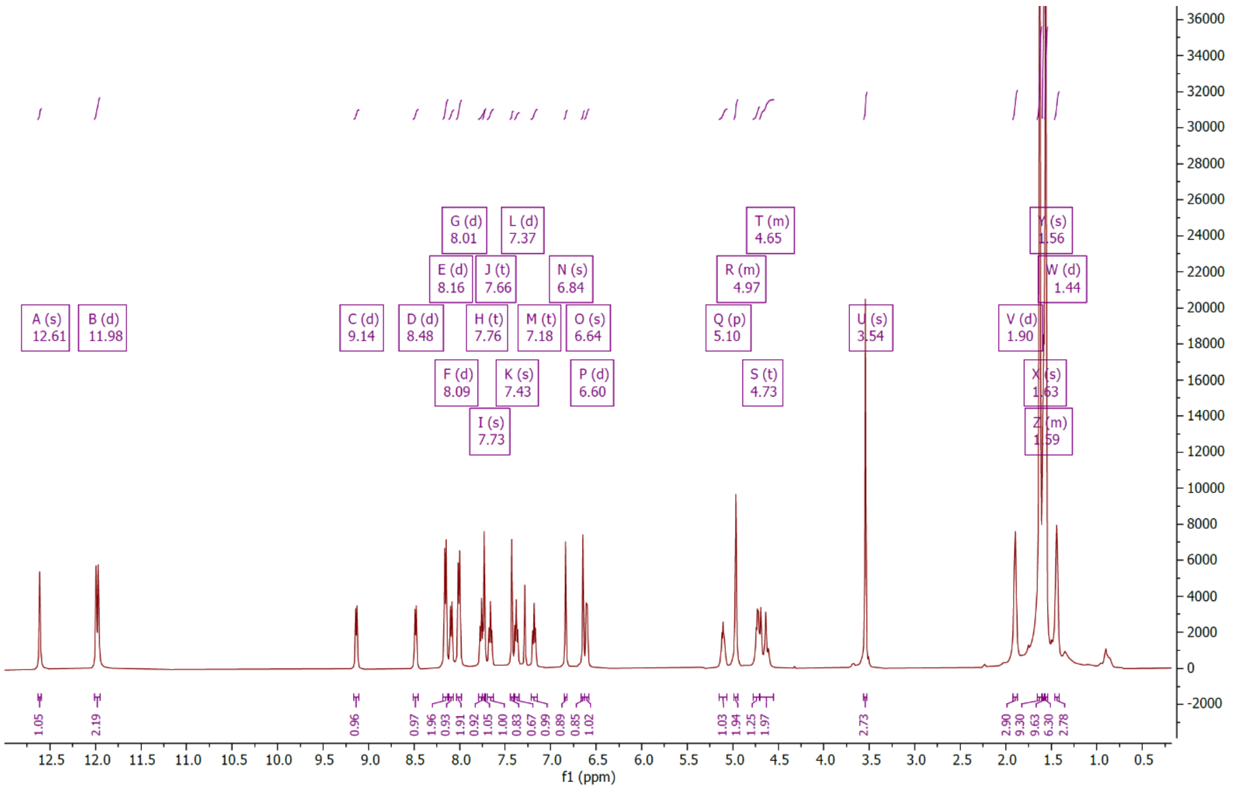

**Synthesis of SK-129$_F$**

To a solution of SK-129-NCS (25 mg, 0.022 mmol) in pyridine (5 ml, anhydrous), N, N-diisopropylethylamine (0.011 ml, 0.066 mmol) was added and the solution was stirred for 10 min. in the atmosphere of argon (g). To this solution, 5-(aminoacetamido) fluorescein (13.5 mg, 0.033 mmol) was added and the reaction was started in dark overnight with continuous stirring in the atmosphere of argon (g). The reaction solution was dried on rotovap under vacuum and the product was purified using column chromatography (0-20% methanol in dichloromethane with 1% triethylamine, v/v) as an orange solid (26.7 mg, 79%).The compound (*tert*-butyl SK-129$_F$) was used in the next step without further characterization. To a solution of *tert*-butyl SK-129$_F$ (20 mg, 0.013 mmol) in dichloromethane (3 mL), triethylsilane (0.1 mL) was added, followed by the addition of trifluoroacetic acid (0.3 mL) and the reaction solution was stirred in dark at room temperature for 4 h. The solution was dried on rotovap in dark and the orange solid was washed with cold diethyl ether (3×5mL), which afforded the desired product (SK-129$_F$) as an orange solid (14.3 mg, 77%). The compound was redissolved in DMSO and purified using HPLC with buffer A (95% water, 5% acetonitrile, 0.1% TFA) and buffer B (95% acetonitrile, 5% water, 0.1% TFA). The gradient for buffer A to B was used from 100% to 0% for a total time of 20 min. at a rate of 3 mL/min. on a reverse-phase C-18 semiprep column (Hypersil gold, 150 mm × 10 mm). The retention peak for SK-129$_F$ was observed around 11 min. $^1$H NMR (500 MHz, DMSO) δ 11.98 – 11.92 (s, 1H), 11.91 – 11.85 (s, 1H), 11.41 – 11.35 (s, 1H), 11.29 – 11.23 (t, *J* = 6.0 Hz, 3H), 10.24 – 10.12 (s, 1H), 9.65 – 9.57 (s, 1H), 8.62 – 8.47 (m, 2H), 8.09 – 8.05 (d, *J* = 7.0 Hz, 1H), 8.04 – 8.00 (d, *J* = 7.6 Hz, 1H), 7.70 – 7.61 (m, 1H), 7.60 – 7.54 (d, *J* = 7.6 Hz, 2H), 7.52 – 7.45 (m, 2H), 7.39 – 7.32 (m, 3H), 7.30 – 7.24 (dt, *J* = 8.0, 4.1 Hz, 1H), 7.15 – 7.05 (d, *J* = 8.7 Hz, 2H), 6.95 – 6.86 (dd, *J* = 11.1, 6.8 Hz, 2H), 6.84 – 6.78 (d, *J* = 7.2 Hz, 1H), 6.65 – 6.56 (m, 1H), 6.48 – 6.42 (m, 1H), 6.31 – 6.09 (m, 7H), 4.99 – 4.80 (m, 4H), 4.79 – 4.67 (dt, *J* = 12.6, 7.0 Hz, 1H), 4.64 – 4.48 (m, 2H), 4.47 – 4.34 (dt, *J* = 11.9, 5.9 Hz, 1H), 1.18 – 1.12 (m, 12H). HRMS-ESI (m/z): calculated for $C_{74}H_{59}N_{10}O_{19}S^+$[(M+H)$^+$]: 1423.3679, found 1423.3684. Anal. Calcd for $C_{74}H_{58}N_{10}O_{19}S$: C, 62.44; H, 4.11; N, 9.84; O, 21.36. Found: C, 62.08; H, 4.38; N, 9.98.

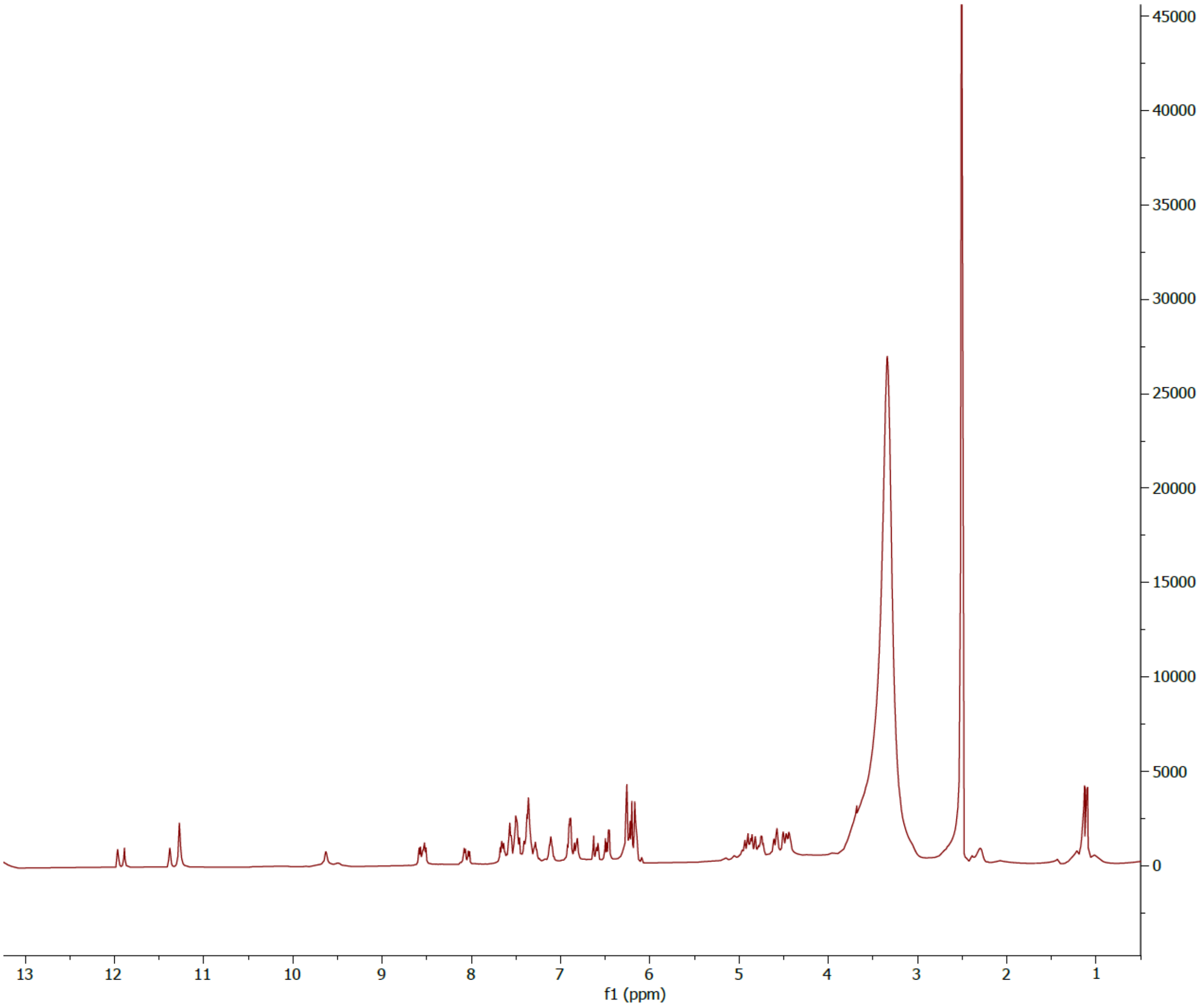


**$^1$H NMR of SK-129$_F$**

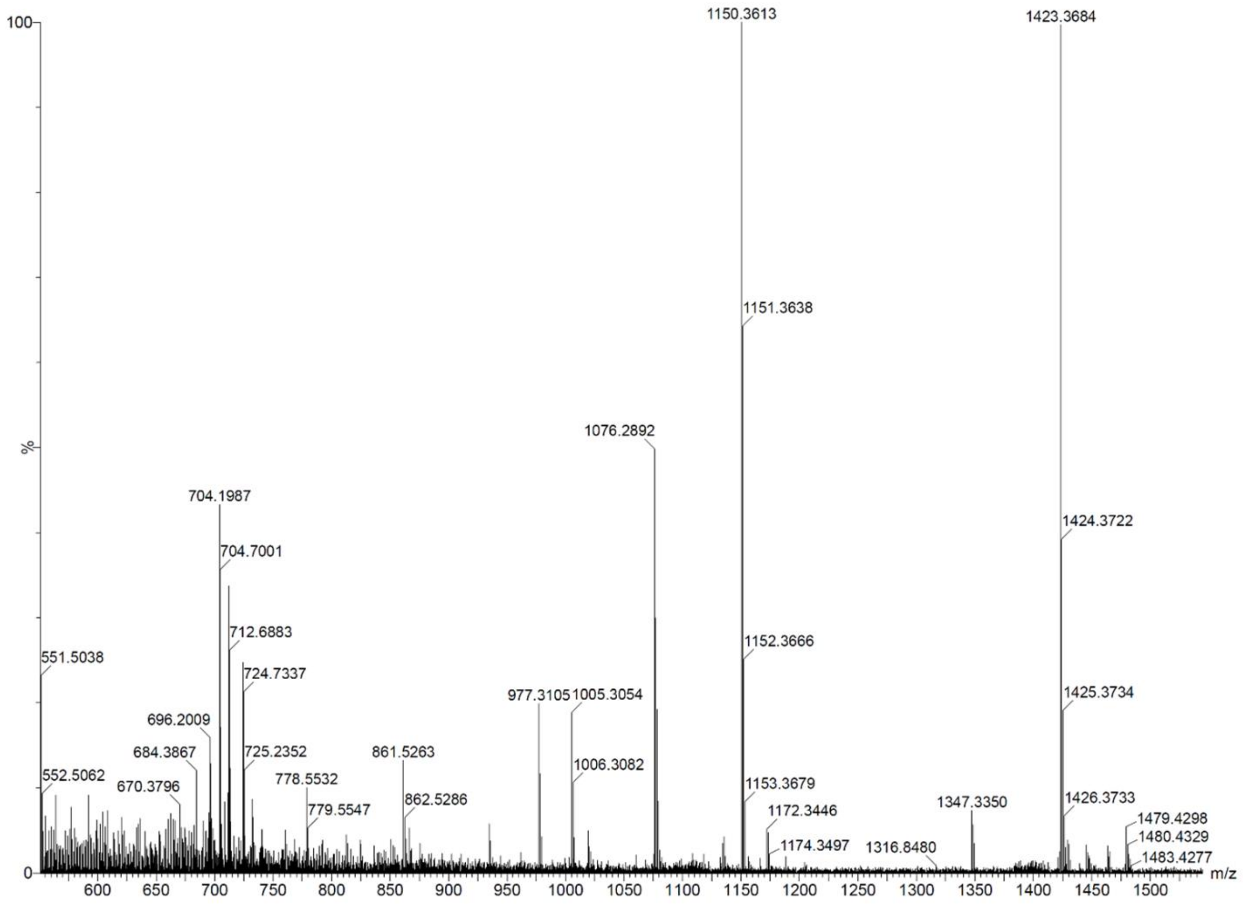


**ESI-HRMS of SK-129$_F$**

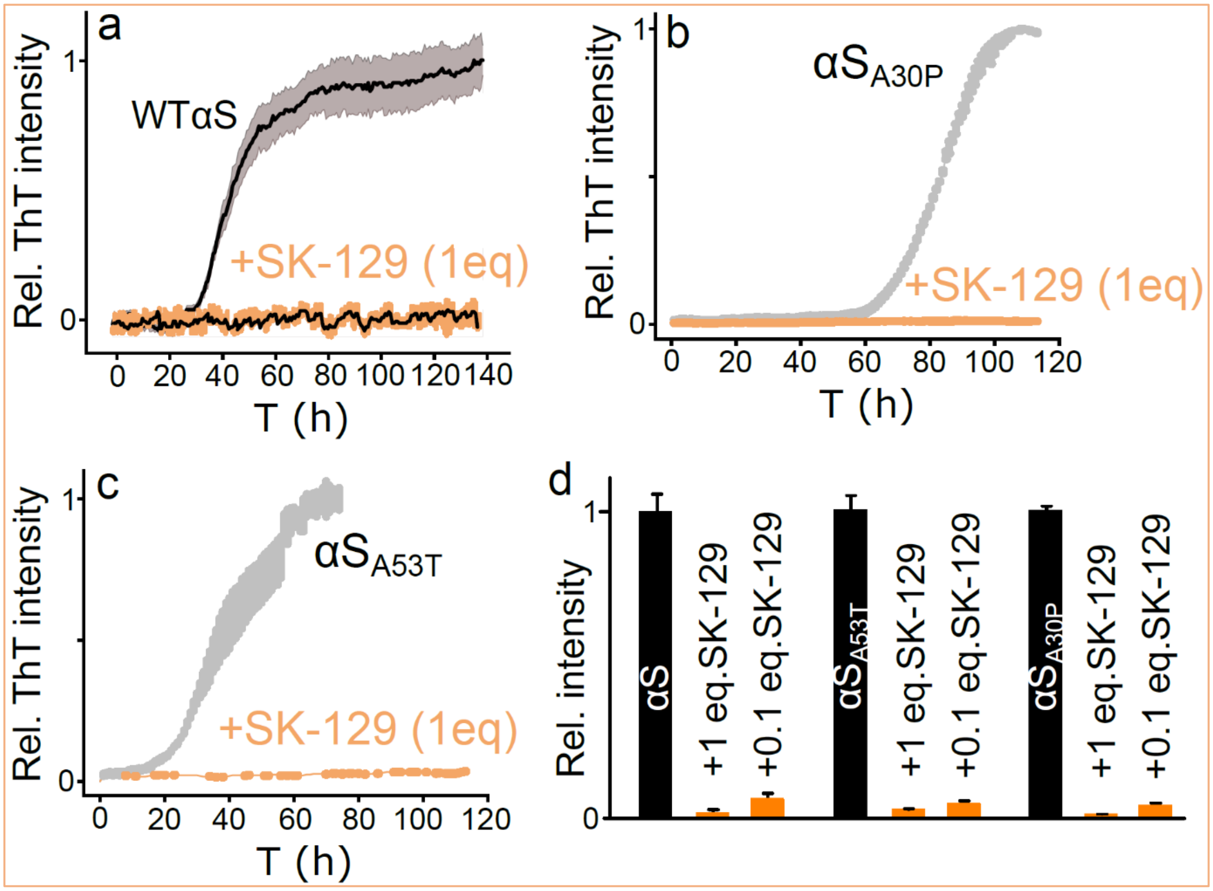


**Fig. 1**. The effect of SK-129 on the aggregation profile of various mutants of αS. The aggregation kinetic profiles of 100 μM WT αS (**a**), 70 μM $\alpha S_{A30P}$ (**b**), and 70 μM $\alpha S_{A53T}$ (**c**) in the absence (light **black curve**) and presence (**Orange curve**) of SK-129 at an equimolar ratio. The aggregation profiles of various αS variants in the absence and the presence of SK-129 are a combination of three different trials. **d**, Statistical analysis of the ThT intensity change for various αS variants in the absence and presence of SK-129 at the indicated stoichiometric ratios. The ThT intensity was measured after 120 h of aggregation of various αS variants in the absence and presence of SK-129. The aggregation kinetics were conducted three times and the reported change in ThT intensity for various conditions is an average of three separate experiments and the error bars are the s.d.'s for three sets of experiments.

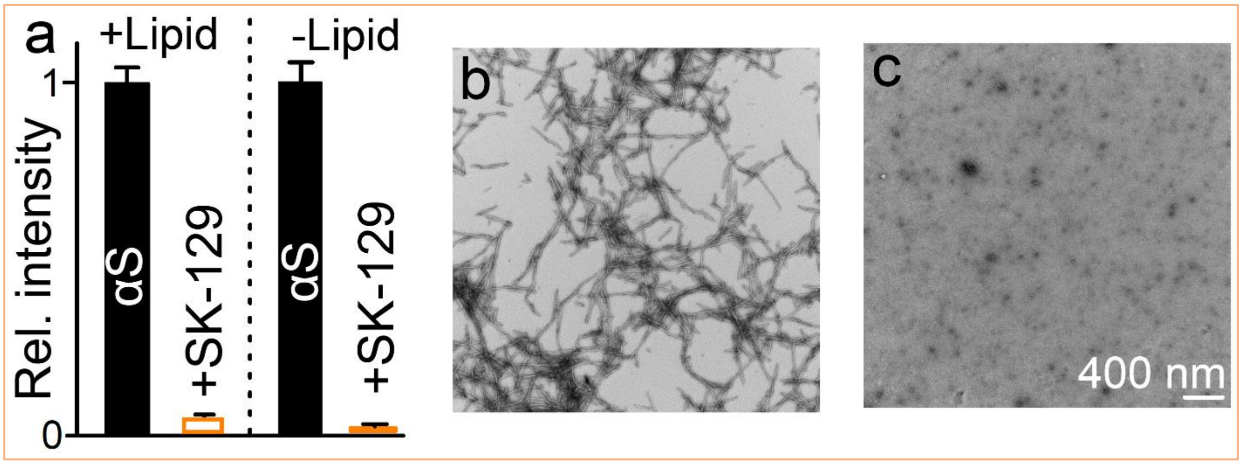


**Fig. 2. a**, The graphical representation of the ThT intensity of the aggregation of 70 μM αS and 35 μM αS under *de novo* conditions and in the presence of LUVs (875 μM, 100 nm, DOPS), respectively in the presence of SK-129 at an equimolar ratio. The ThT intensity of the lipid-free and lipid-catalyzed aggregation of αS was monitored after 7 days. Negatively stained-TEM images of the aggregation of 70 μM αS in the absence (**b**) and presence (**c**) of SK-129 at an equimolar ratio after four days. The aggregation kinetics were conducted three times and the reported change in ThT intensity for various conditions is an average of three separate experiments. The reported error bars are the s.d.'s for multiple sets of experiments.

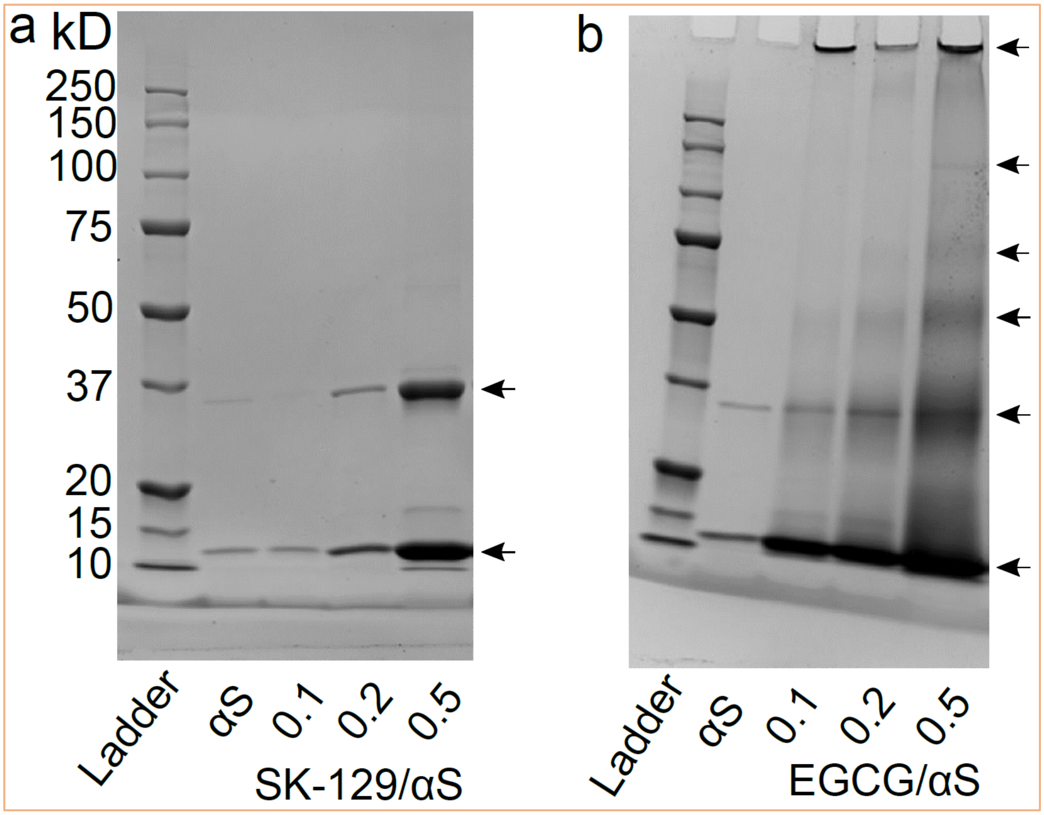


**Fig. 3**. Gel-shift images of αS (70 µM) incubated in the absence (lane 2) and presence of SK-129 (**a**) and EGCG (**b**) at sub-stoichiometric ratios (0.1, 0.2, 0.5 mol eq.) for 7 days at 37$^{0}$C with constant shaking. The solutions of αS in the absence and presence of ligands were centrifuged and the supernatant was used for the gel shift assay. The arrows indicate the formation of various αS structures. The gel shift assay of the supernatant of the αS aggregated solution resulted in a very small amount of monomer because most of the protein converted into fibers. Most of the αS was found in the supernatant as a monomer and a dimer in the presence of SK-129; however, higher-order oligomers were observed in the presence of EGCG. The most toxic states of αS are considered to be the higher-order oligomers (n>5) and SK-129 efficiently inhibits their formation.

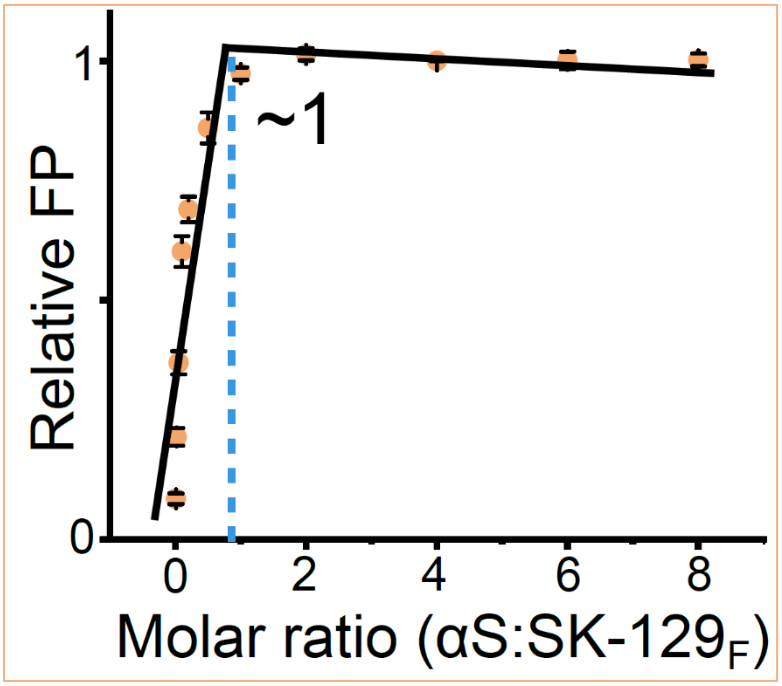


**Fig. 4**. The plot between the relative FP and the molar ratio (αS:SK-$129_F$), which is extracted from the FP titration between 10 μM SK-$129_F$ and increasing concentrations of αS. The binding stoichiometry of SK-$129_F$ against αS was determined by fit using two linear equations. The intersection of two equations lead to the binding stoichiometric ratio between αS and SK-$129_F$. The fluorescence polarization titrations between $SK129_F$ and αS were conducted at least three times and each point in titrations was the average of three data points. The reported error bars are the s.d.'s for multiple sets of experiments conducted on separate occasions.

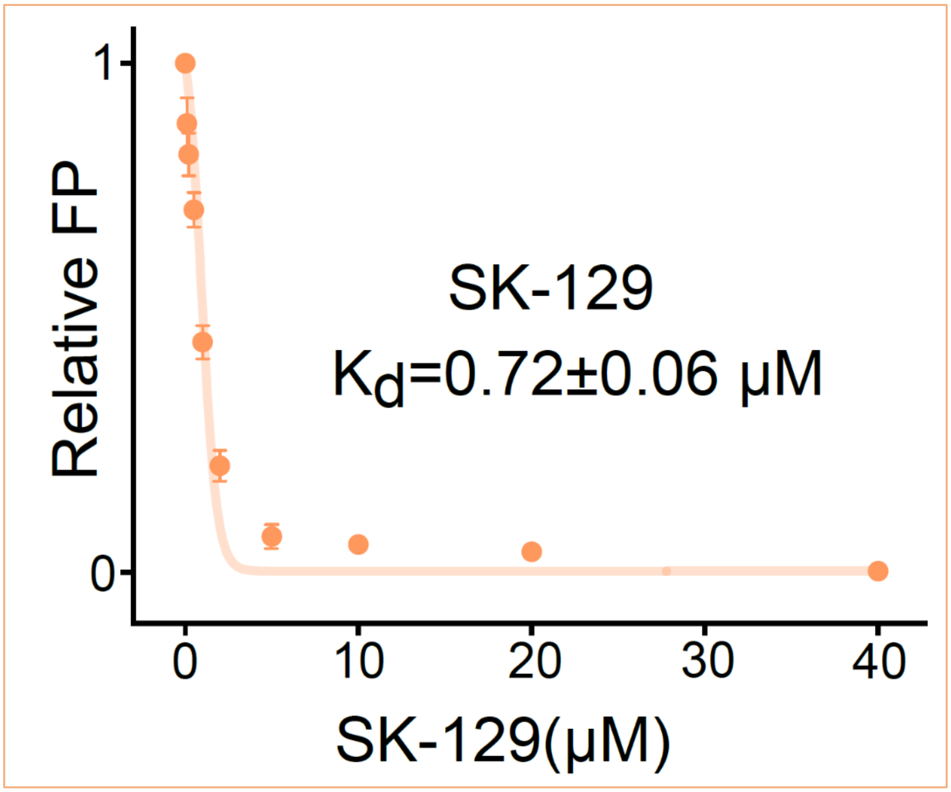


**Fig. 5**. A competitive FP titration between a preformed saturated solution of αS-SK-129$_F$ (100 μM: 10 μM) and SK-129 to determine the binding affinity between SK-129 and αS. SK-129 was serially added to a saturated solution of αS-SK-129$_F$ until no more change in the FP was observed. The plot between the related change in the FP against the concentration of SK-129 was fit using a competitive one binding site model to determine the binding affinity between SK-129 and αS. The $K_d$'s for SK-129 and SK-129$_F$ against αS were 0.72±0.06 and 0.80±0.06, respectively, which suggests that the fluorescein tag on SK-129$_F$ has a slight effect on the binding affinity of SK-129 against αS. The fluorescence polarization titrations between SK129 and the preformed saturated complex of SK129$_F$-αS were conducted at least three times and each point in titrations was the average of three data points. The reported error bars are the s.d.'s for multiple sets of experiments conducted on separate occasions.

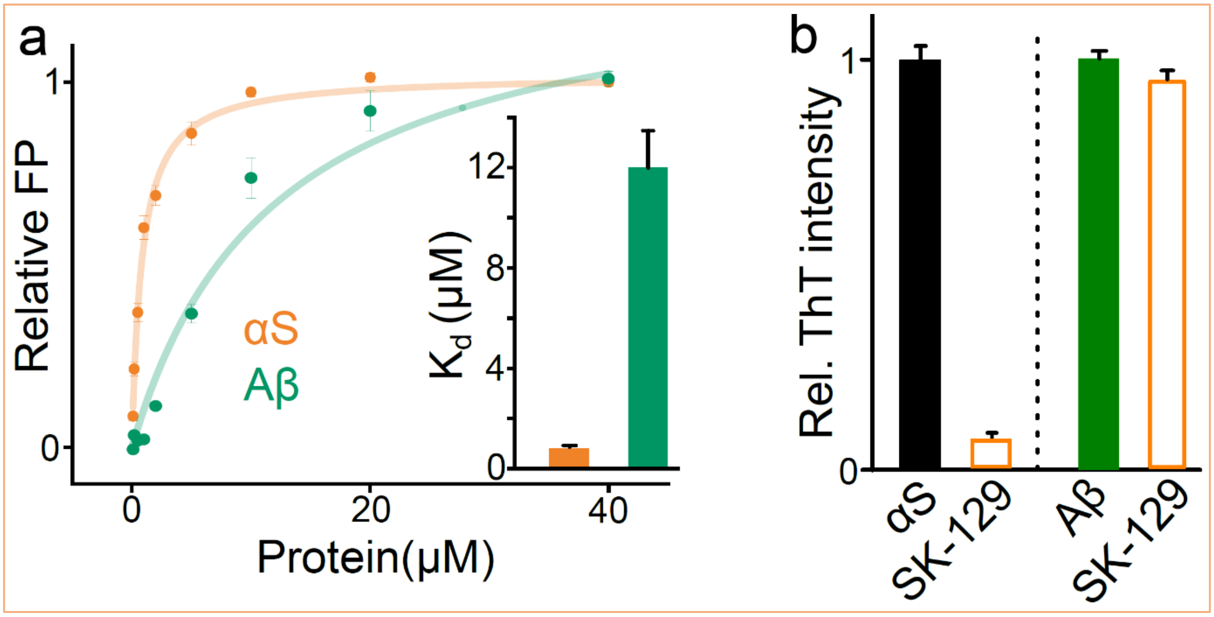


**Fig. 6**. **a,** The curves and graphical representation (inset) of the comparison of FP-based binding affinities of SK-129F against αS and Aβ. **b**, The comparison of the antagonist activity of SK-129 against the aggregation of 70 μM αS and 15 μM Aβ at an equimolar ratio. The final ThT intensity was monitored after one and four days for Aβ and αS, respectively. The fluorescence polarization titrations between SK129 and various proteins were conducted at least three times and each point in titrations was the average of three data points. The reported error bars are the s.d.'s for three sets of experiments.

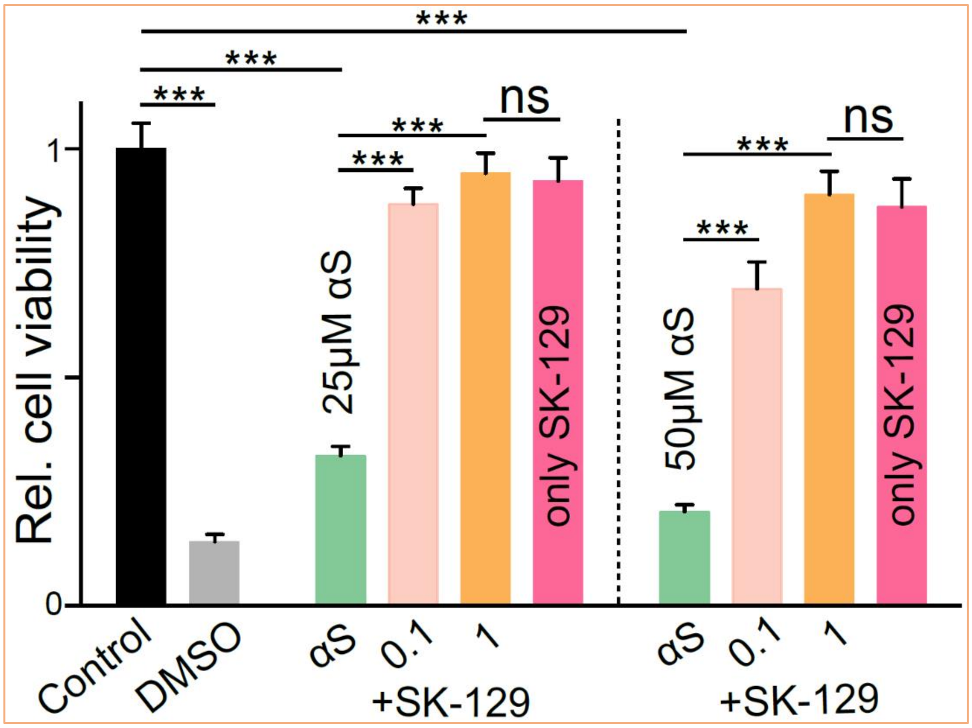


**Fig. 7**. The statistical analysis of the relative viability of SH-SY5Y cells in the presence of the indicated concentrations of αS and the αS-SK-129 complex (the aggregated state) at the indicated molar ratios using the MTT assay. For the relative viability the highest and lowest cell viability are used from the control (1 × PBS buffer) and DMSO, respectively. The inherent toxicity of SK-129 in SH-SY5Y was also monitored at the indicated concentrations. The cell viability assays were conducted with at least four biological replicates and four technical replicates for each biological replicate and the reported error bars are the s.d.'s for each experiment. Statistical significance was analyzed using a one-way analysis of variance (ANOVA) with Tukey's multiple comparison's test. *p<0.05, **p<0.01, ***p<0.001.

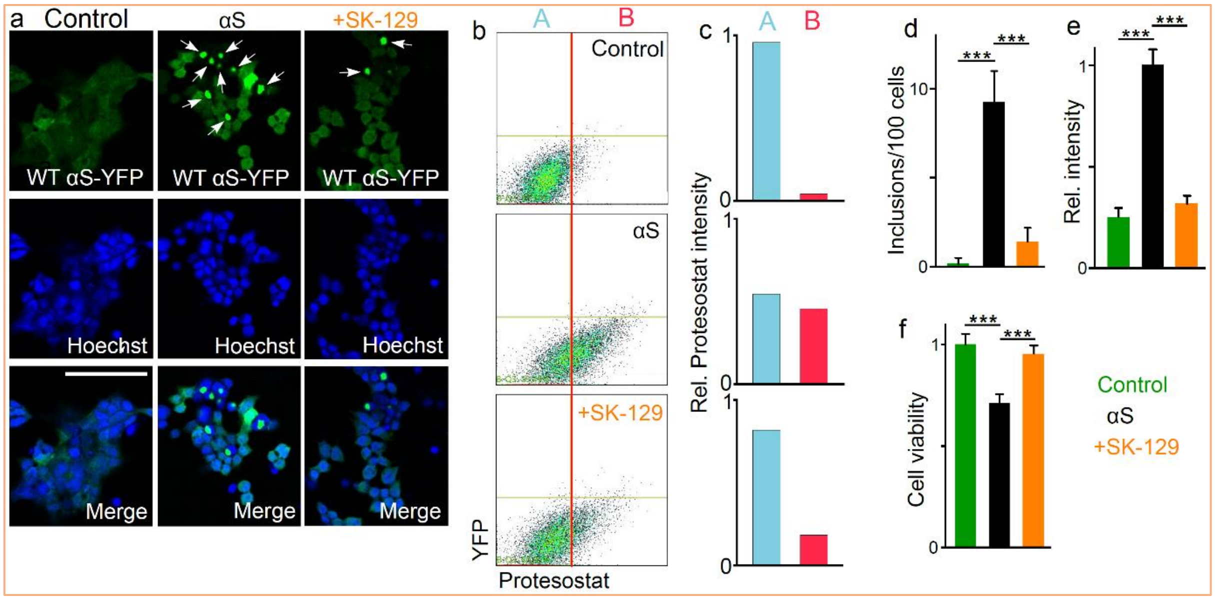


**Fig. 8**. The effect of αS fibers on the HEK cells (αS-YFP) in the absence and presence of SK-129. **a**, The confocal images of the HEK cells (expressing endogenous WT αS-YFP) treated with the control (1 × PBS buffer), 7 μM αS (the aggregated state), and 7 μM αS-SK-129 complex (the aggregated state). The WT αS-YFP inclusions are indicated with white arrows. The images show the staining of HEK cells with Hoechst dye (blue) and the merge is the combination of the Hoechst and YFP signals. scale bar, 100 μm. **b**, The Flow cytometry analysis of HEK cells treated with the indicated conditions. The x-axis represents αS-YFP aggregates containing cells stained with Proteostat dye and the Y-axis is for the YFP signal intensity. **c**, A and B represent the relative % of HEK cells without and with αS-YFP aggregates, respectively. **d**, The number of inclusions in HEK cells observed for the indicated conditions. Relative intensity of ProteoStat-stained aggregates (**m**) and relative viability (**n**) of HEK cells under the indicated conditions. The cell viability assays and the protesostat intensity measurement were conducted with at least four biological replicates and four technical replicates for each biological replicate and the reported error bars are the s.d.'s for each experiment. The counting of inclusions was carried out for six different experiments and for each experiment 100 cells were counted from at least four different locations. Statistical significance was analyzed using a one-way analysis of variance (ANOVA) with Tukey's multiple comparison's test. *p<0.05, **p<0.01, ***p<0.001.

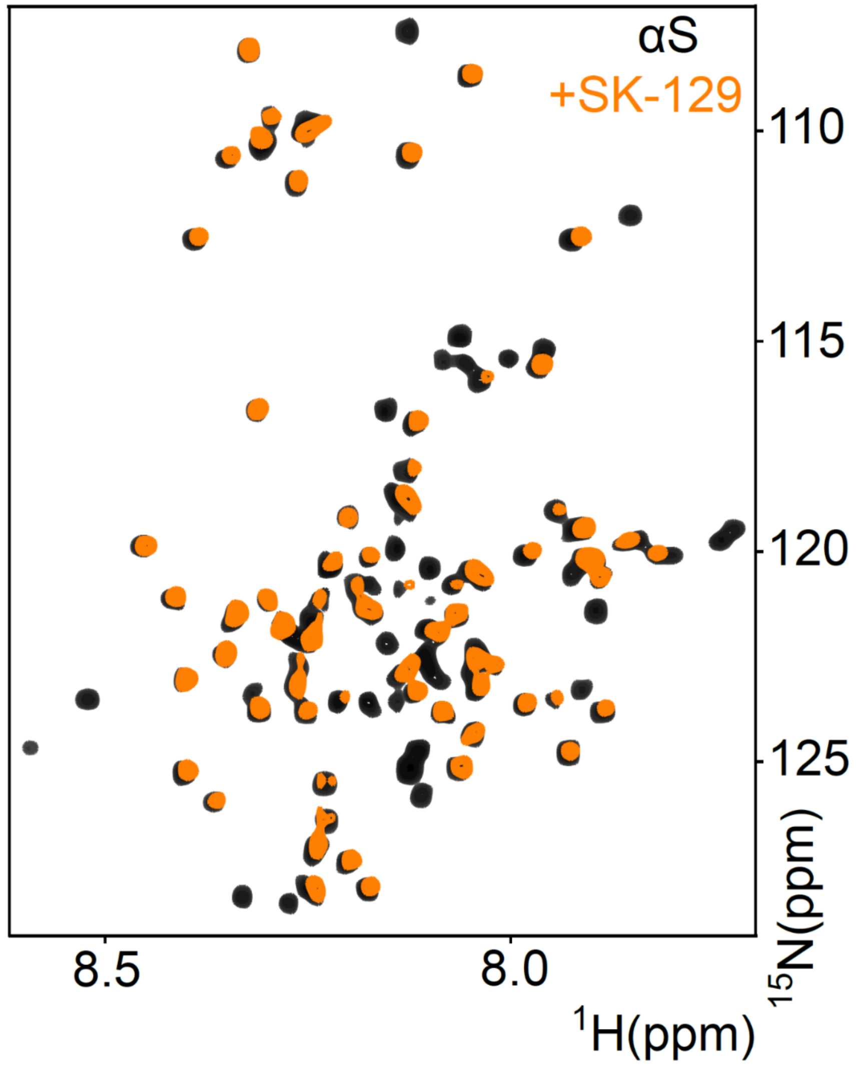


**Fig. 9**. Overlay of two-dimensional HSQC ($^{1}$H, $^{15}$N) NMR spectra of 70 µM uniformly $^{15}$N-labelled αS in the absence (black) and presence (yellow) of 140 µM SK-129. The HSQC NMR experiment conditions were exactly similar to the HSQC NMR spectrum at an equimolar ratio (SK-129: αS) (Main Manuscript Figure 2).

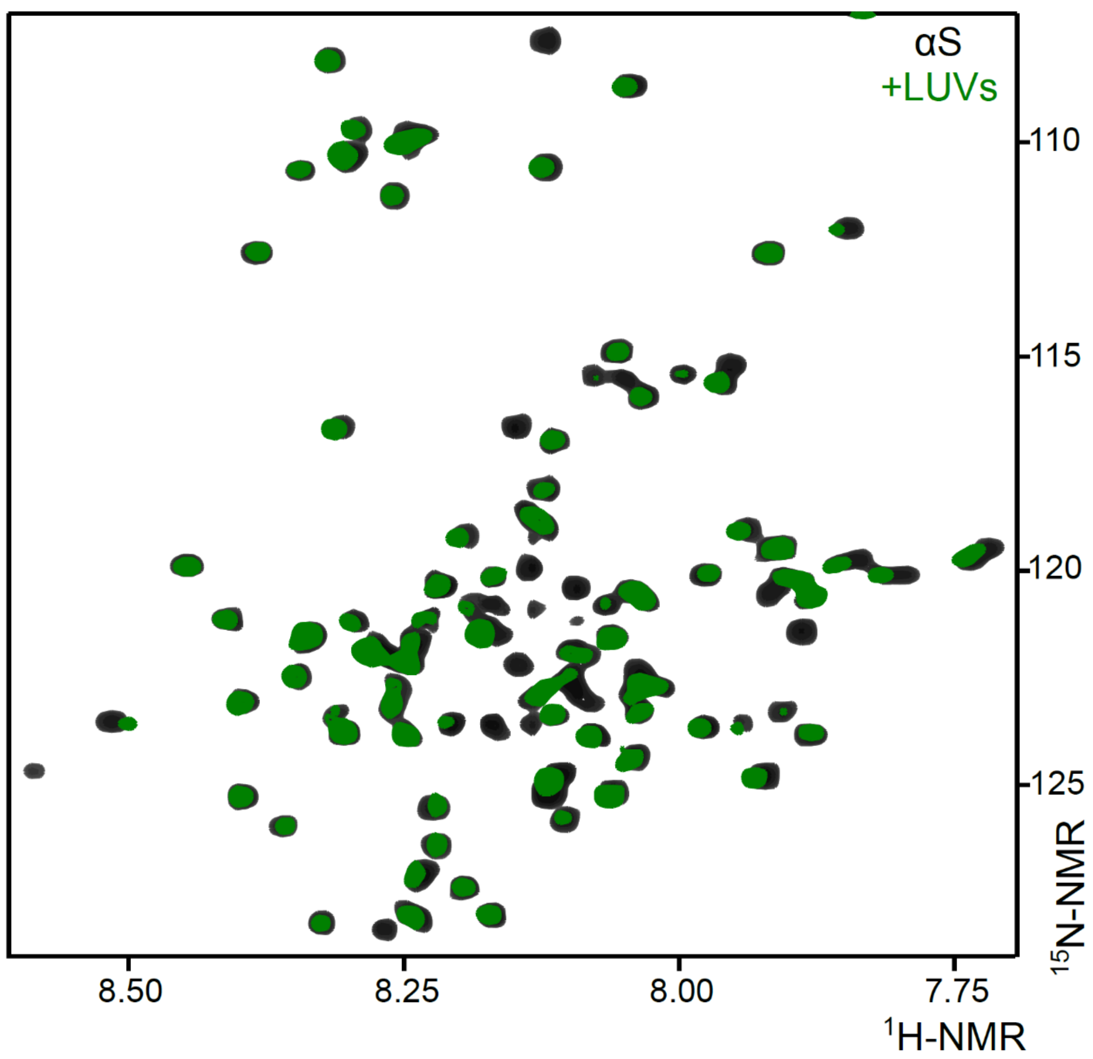


**Fig. 10**. Overlay of two-dimensional HSQC ($^1$H, $^{15}$N) NMR spectra of 70 μM uniformly $^{15}$N-labelled αS in the absence (black) and presence (yellow) of 875 μM LUVs (100 nm, DOPS). The HSQC NMR experiment conditions were exactly similar to the HSQC NMR spectrum at an equimolar ratio (SK-129: αS) (Main manuscript Figure 2).

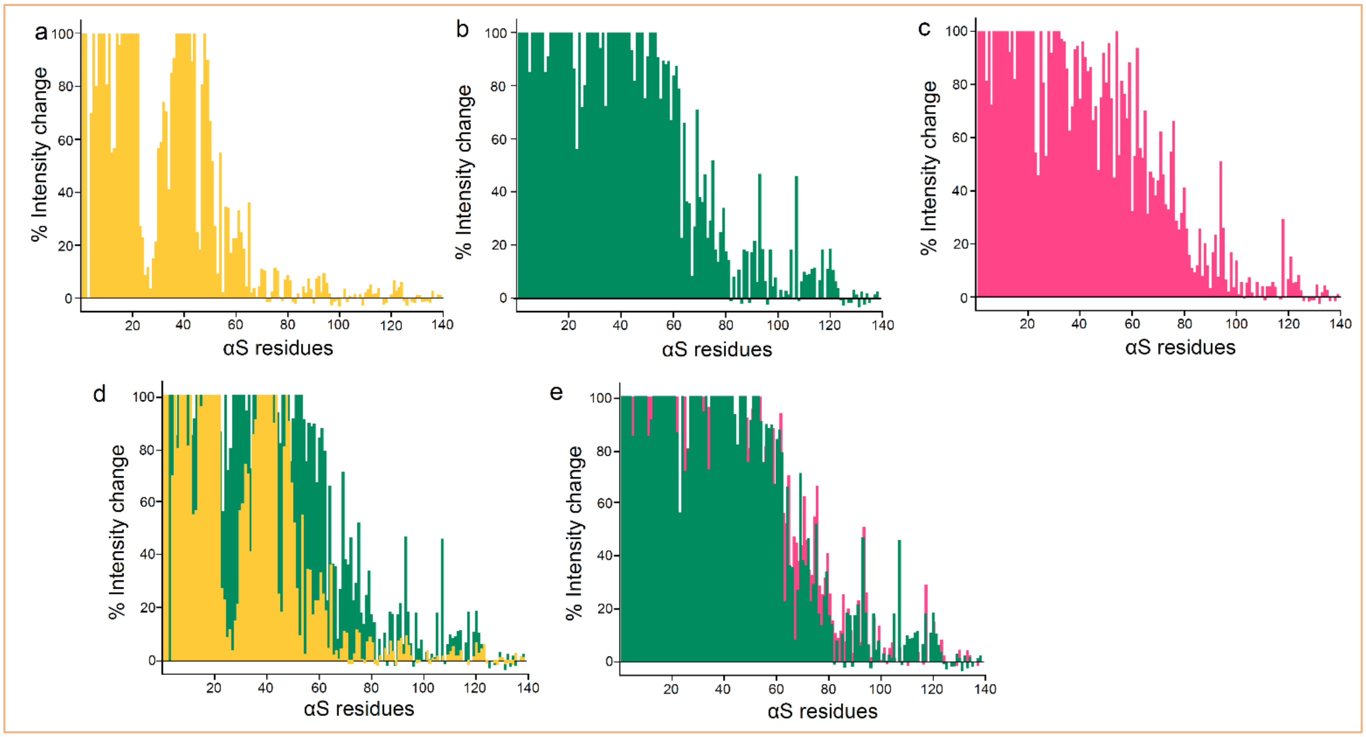


**Fig. 11**. Comparison of the binding interaction between SK-129 and αS, and αS and LUVs using HSQC NMR spectroscopy. Graphical presentation of the intensity changes of the backbone amide peaks of $^{15}$N-labelled αS (70 µM) in the presence of 70 µM (**a**) and 140 µM (**b**) SK-129, and LUVs (875 µM, 100 nm, DOPS) (**c**). **d**, Overlay of the intensity changes of the backbone amide peaks of $^{15}$N-labelled αS (70 µM) in the presence of 70 µM (orange) and 140 µM (green) SK-129. **e**, Overlay of the intensity changes of the backbone amide peaks of $^{15}$N-labelled αS (70 µM) in the presence of 140 µM SK-129 (green) and 875 µM LUVs (red).

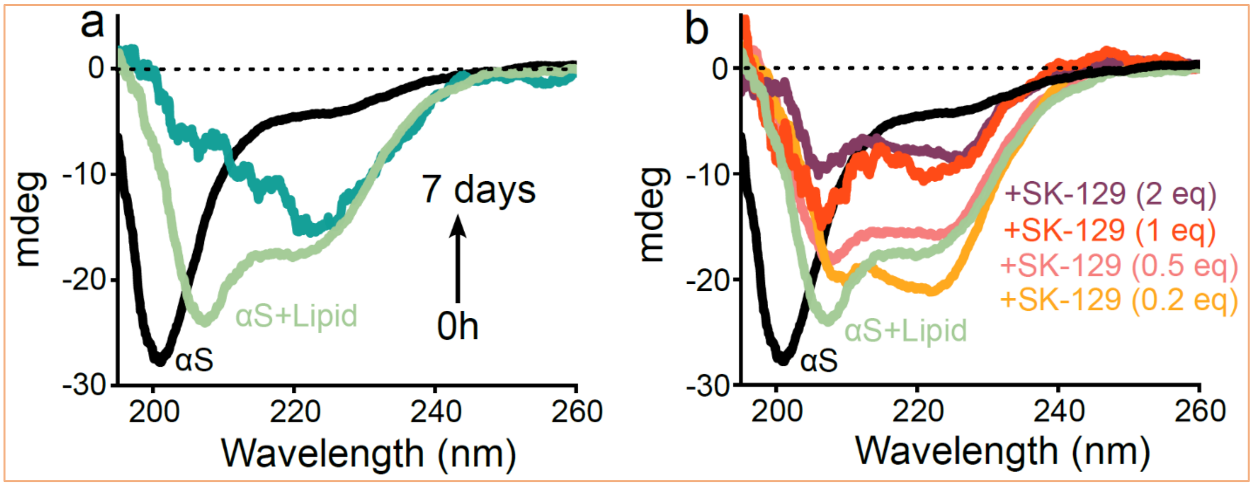


**Fig. 12**. CD-based characterization of the effect of SK-129 on lipid-catalyzed aggregation kinetics of αS. **a**, Time-dependent CD spectra of 30 µM αS in the absence (black) and presence of LUVs (375 µM, 100 nm, DOPS) for 7 days. **b**, CD spectra of 30 µM αS in the presence of LUVs (375 µM, 100 nm, DOPS) and SK-129 under indicated stoichiometric ratios after 7 days.

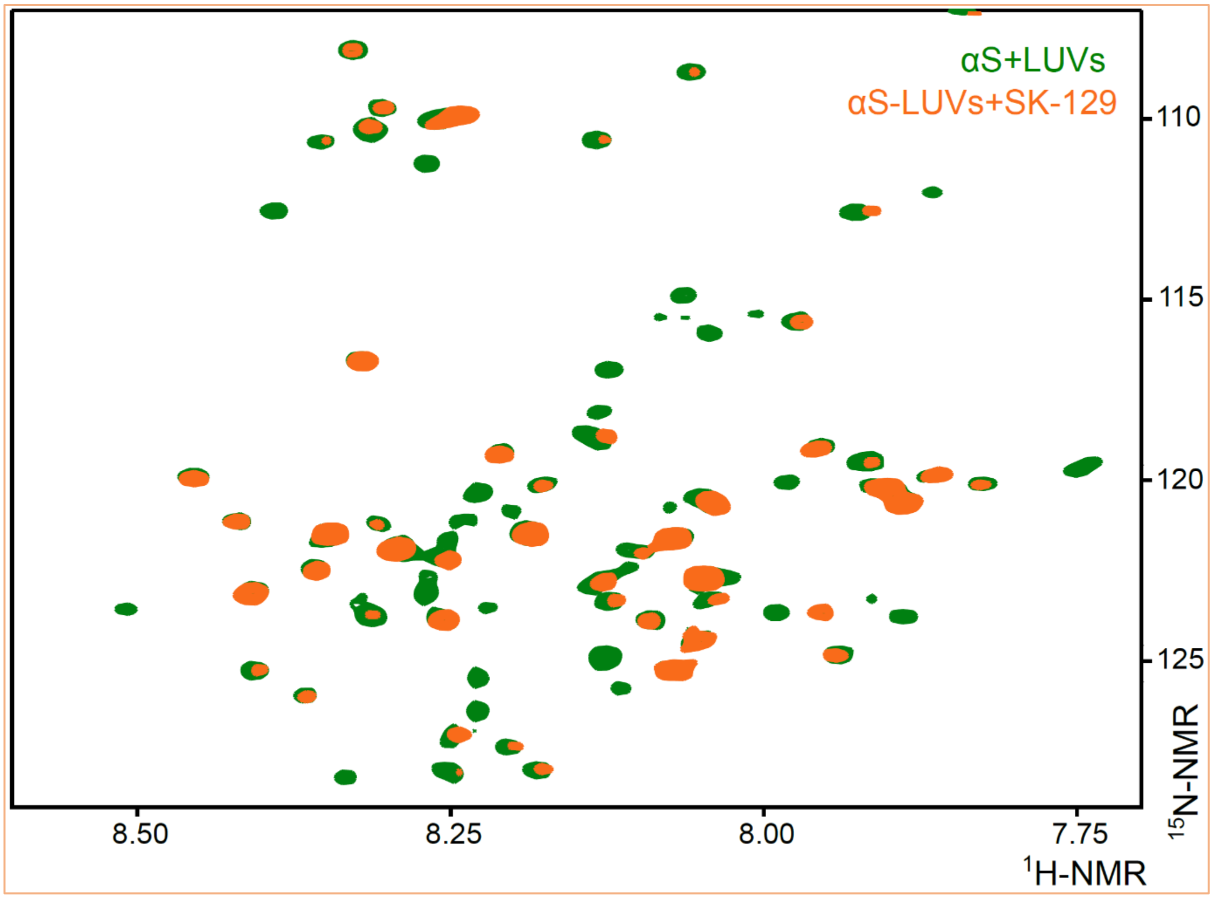


**Fig. 13**. Overlay of two-dimensional HSQC ($^{1}$H, $^{15}$N) NMR spectra of 70 µM uniformly $^{15}$N-labelled αS (+LUVs, 875 µM, 100 nm, DOPS) in the absence (green) and presence (orange) of 140 µM SK-129. The HSQC NMR experiment conditions were exactly similar to the HSQC NMR spectrum at an equimolar ratio (SK-129:αS) (Main Manuscript Figure 2).

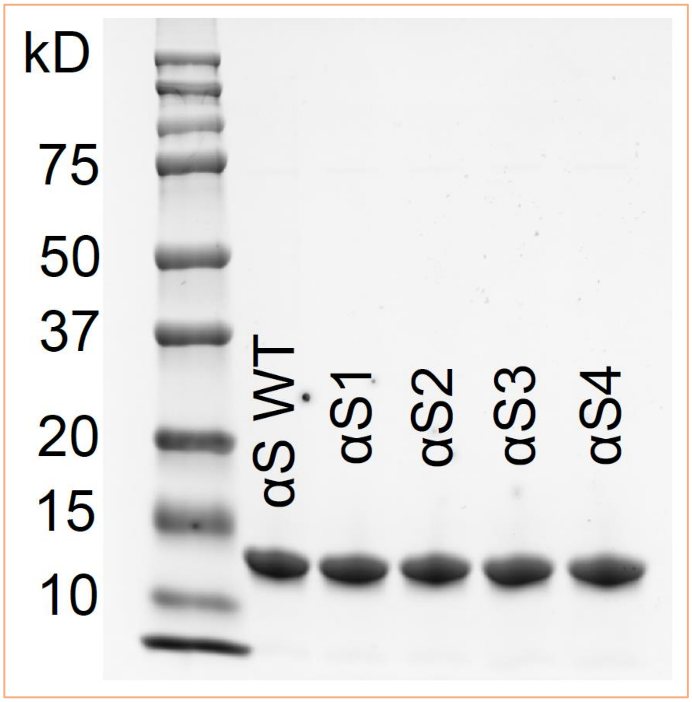


**Fig. 14.** The Gel shift of WT αS and various αS mutants, including αS1, αS2, αS3, and αS4 that have deleted residues 6-12, 15-23, 36-45, and 48-53, respectively.

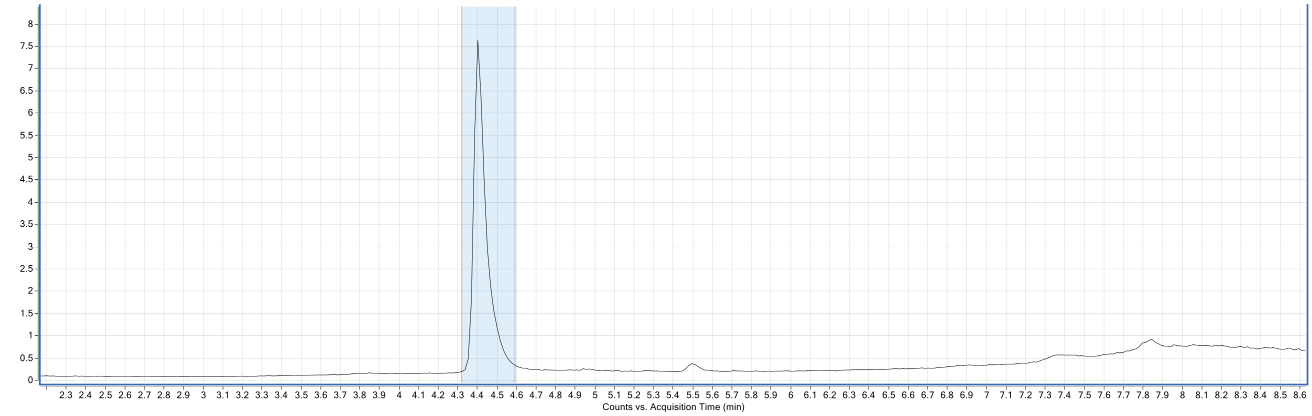


**Fig. 15a**: HPLC chromatogram for WT αS.
Conditions for HPLC run.
**Column**: Agilent PLRP-S 1000A, 5 µm 2.1 mm × 50 mm (PL1912-1502)
**Solvent A**: 100% water in 0.1% Formic Acid
**Solvent B**: 90% aq. Acetonitrile in 0.1% formic acid
**Flow Rate**: 0.3 mL/min
**Gradient**: 0 min – 5% B, 2 min – 5% B, 7 min – 95% B, 9 min – 95% B, 11 min – 5% B, 15 min–stop time.

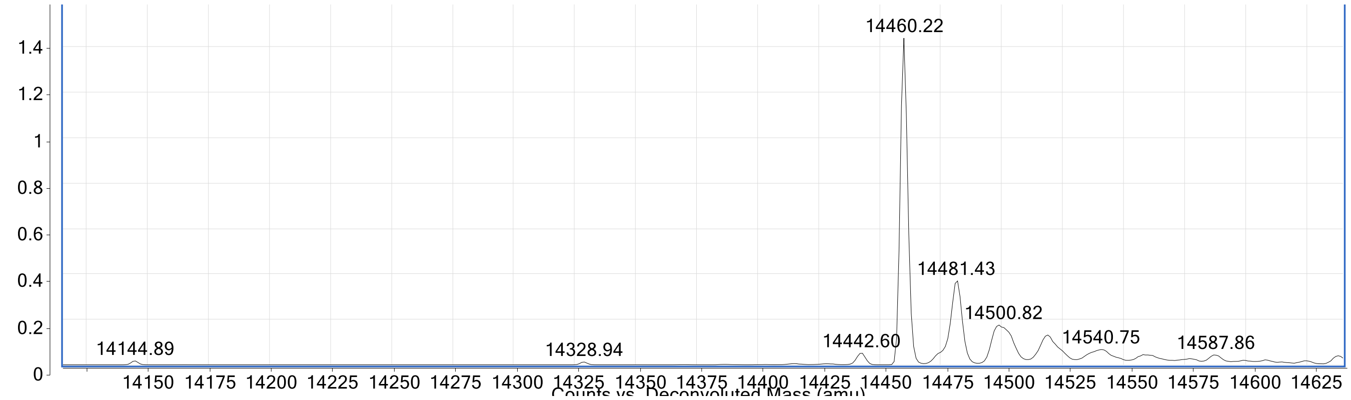


**Fig. 15b.** ESI-MS spectrum of αS1. Theoretical mass = 14460, Observed mass=14460.22.

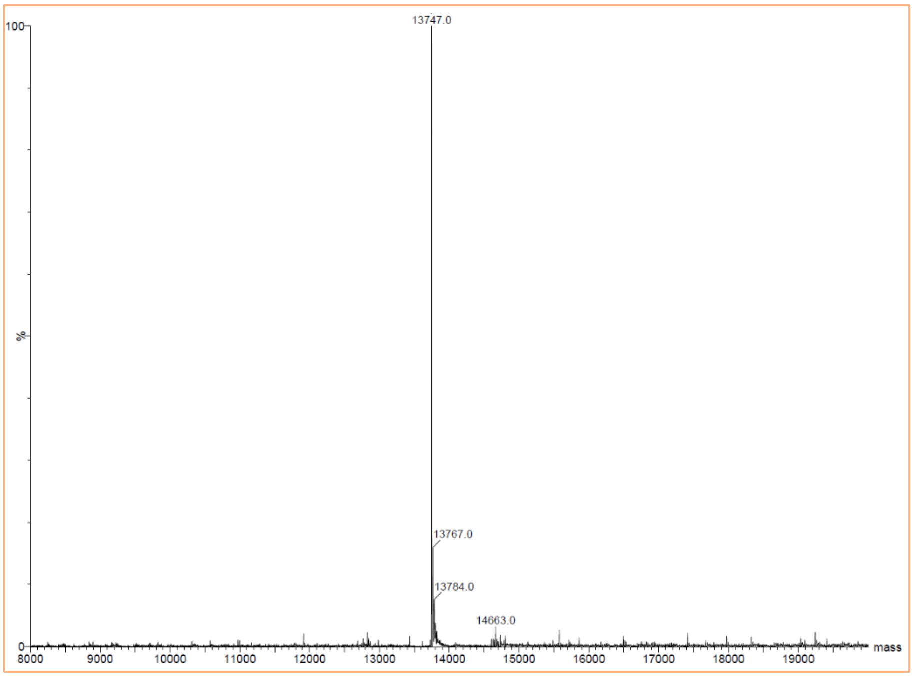


**Fig. 15c.** ESI-MS spectrum of αS1. Theoretical mass = 13,747.27, Observed mass=13,474.0

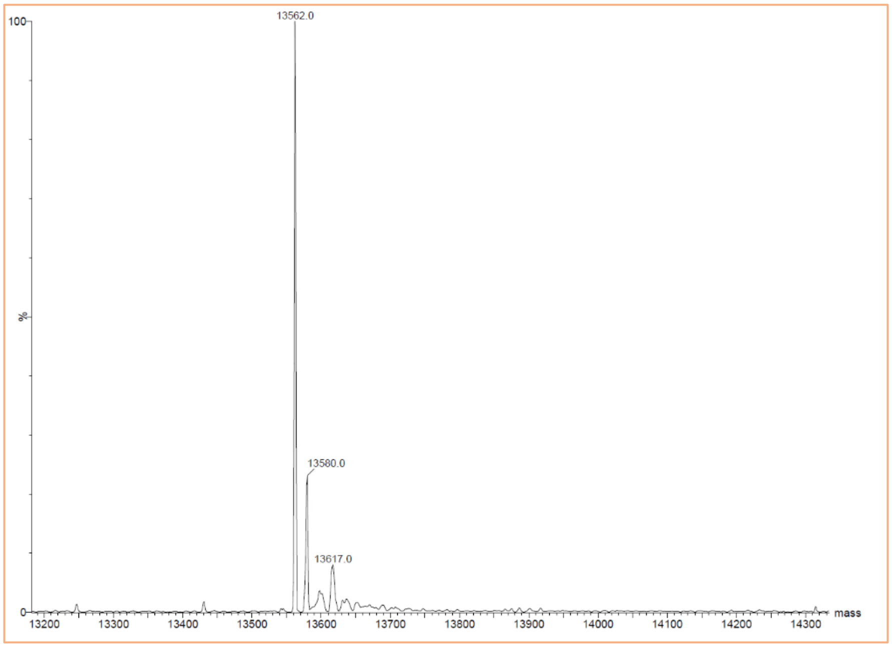


**Fig. 15d.** ESI-MS spectrum of αS2. Theoretical mass = 13,562.09, Observed mass=13,562.0

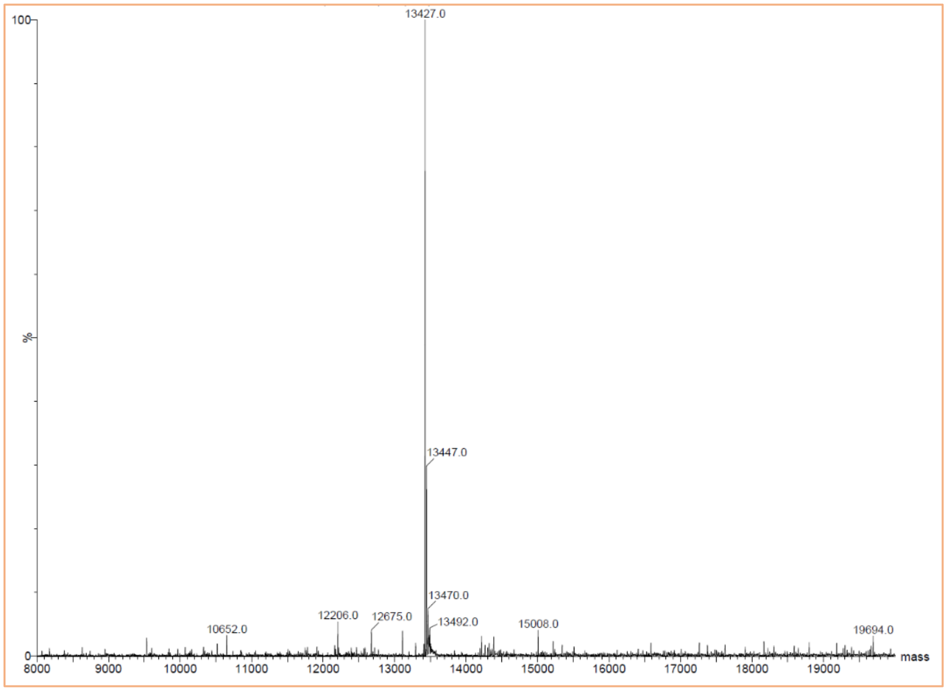


**Fig. 15e.** ESI-MS spectrum of αS3. Theoretical mass = 13,426.92, Observed mass=13,427.0

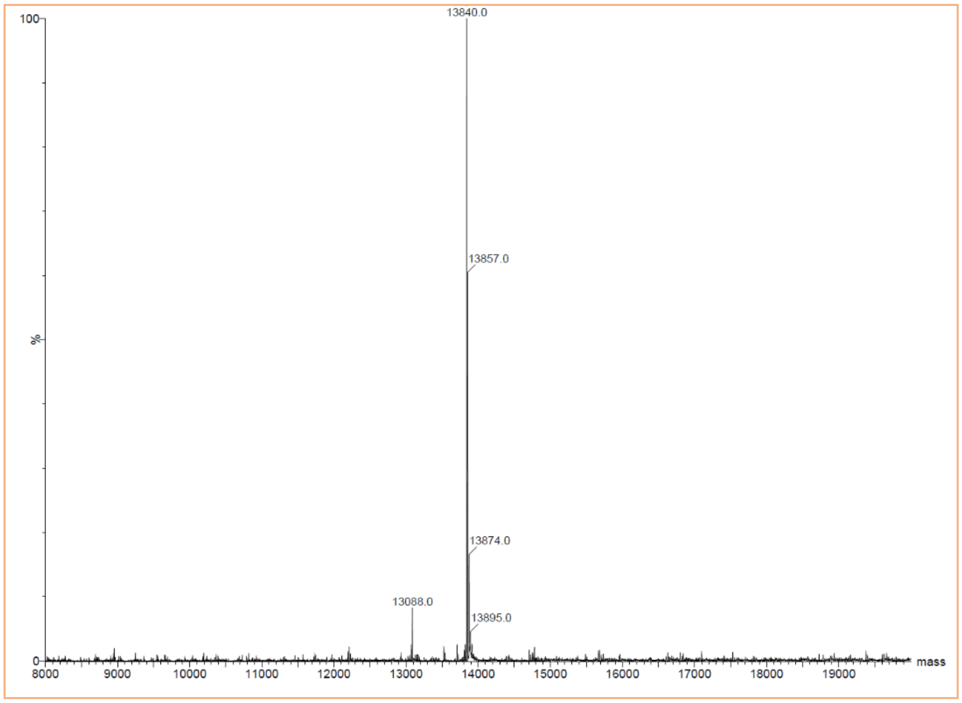


**Fig. 15f.** ESI-MS spectrum of αS4. Theoretical mass = 13,840.43, Observed mass=13,840.0

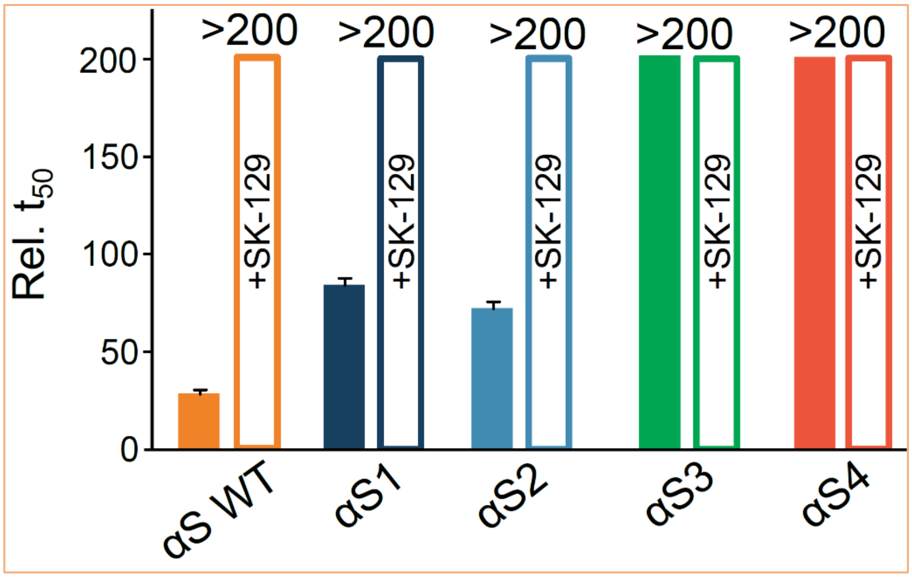


**Fig. 16**. Comparison of $t_{50}$'s (The time required to reach 50% fluorescence intensity of ThT) for the aggregation of 100 μM WT αS and αS mutants in the absence (filled bar) and presence of SK-129 (open bar) at an equimolar ratio. The aggregation kinetics of various proteins were conducted three times and the reported $t_{50}$ for various proteins is an average of three separate experiments. The reported error bars are the s.d.'s for three separate experiments.

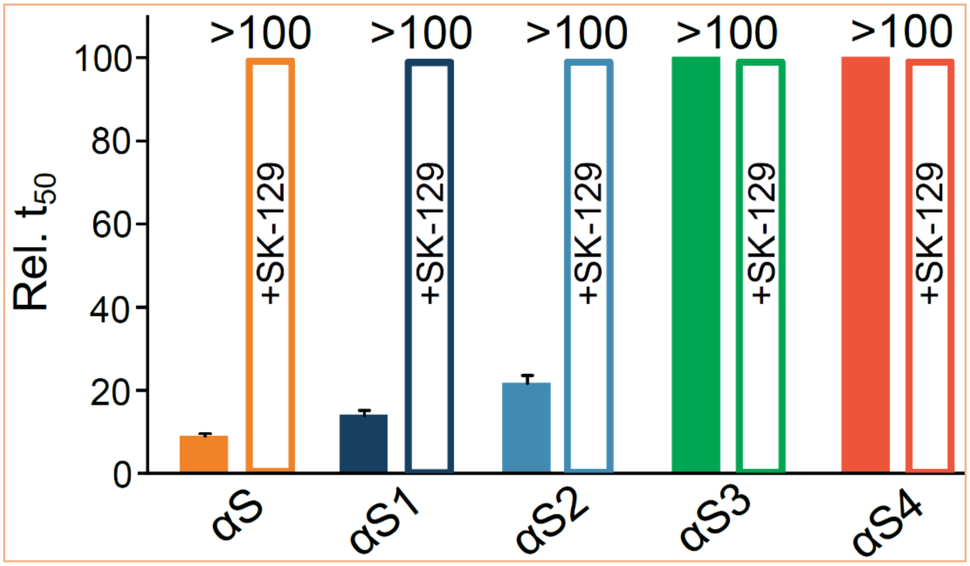


**Fig. 17**. Comparison of $t_{50}$'s (The time required to reach 50% fluorescence intensity of ThT) of seed catalyzed (seeds= WT αS, 10% in monomer concentration of αS) aggregation of 100 μM WT αS and αS mutants in the absence (filled bar) and presence of SK-129 (open bar) at an equimolar ratio. The aggregation kinetics of various proteins were conducted three times and the reported $t_{50}$ for various proteins is an average of three separate experiments. The reported error bars are the s.d.'s for three separate experiments.

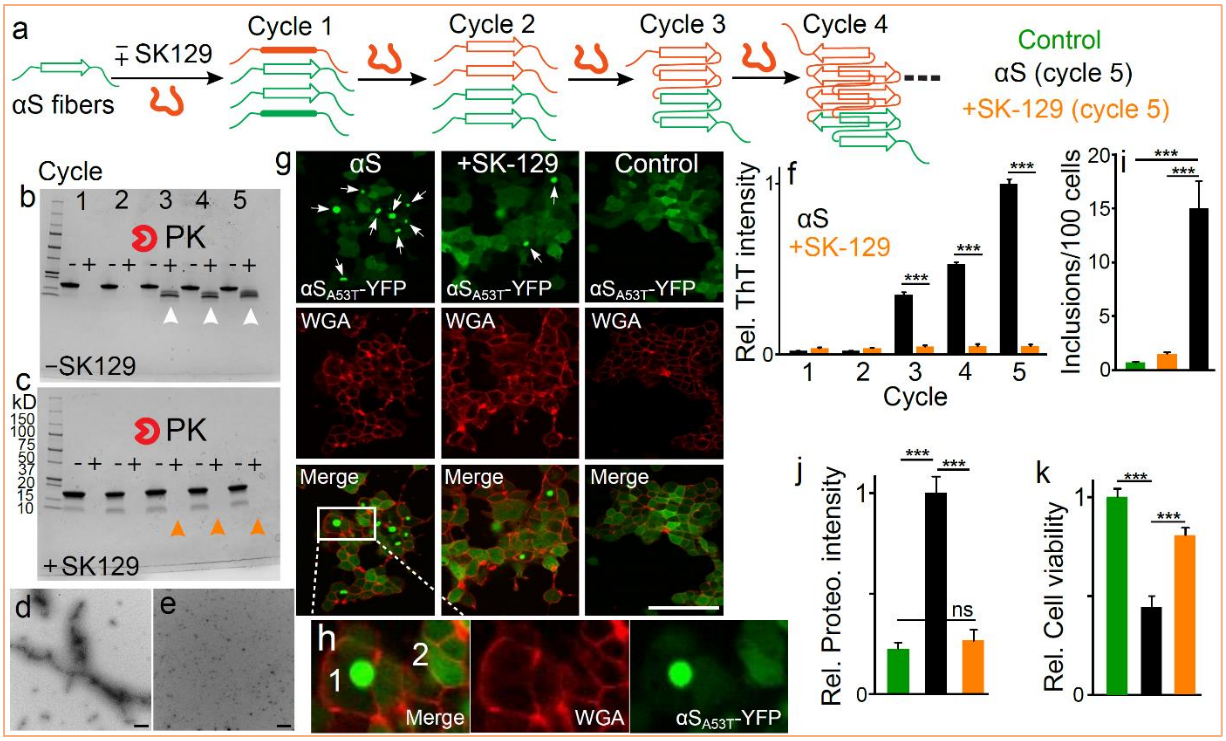


**Fig. 18**. Effect of SK-129 on the prion-like spread of αS. **a**, Schematic of the protein misfolding cyclic amplification (PMCA) assay. In the first step, the preformed fibers of αS (1 µM in monomer conc.) were used to template αS monomer (20 µM) in the absence and presence of 20 µM SK-129. The solutions were incubated at 37 $^0$C with constant shaking for 2 days. In the second step, 1/10 volume of the solutions (±SK-129) was used to template αS monomer. The process was repeated for five cycles. The Bis-tris gels of PMCA samples from cycle first to fifth in the absence (**b**) and presence (**c**) of SK-129. The -ve and +ve signs indicate the amplified samples in the absence and presence of PK, respectively. The arrows indicate the effect of PK on the PMCA samples from the indicated cycles. TEM images of the PMCA samples from the fifth cycle in the absence (**d**) and presence (**e**) of SK-129. Scale bar = 400 nm. **f**, The statistical analysis of the relative ThT intensity of various PMCA samples in the absence (black bar) and presence (orange bar) of SK-129, before treating the samples with PK. The reported change in ThT intensity for various conditions is an average of three separate experiments. The reported error bars are the s.d.'s for three experiments. **g**, The representative images of HEK cells after treatment with PMCA samples from the fifth cycle under the indicated conditions. The $\alpha S_{A53T}$-YFP inclusions are indicated by white arrows. The green color images are due to the intracellularly expressed $\alpha S_{A53T}$-YFP and the red colored images are due to the staining of HEK cell membranes by wheat germ agglutinin (WTA). **h**, The zoom-in version of the difference in an intracellular $\alpha S_{A53T}$-YFP inclusion (1) and the homogenous distribution of αS-$_{A53T}$-YFP (2). Scale bar = 100 µm. **i**, The statistical analysis of the number of αS inclusions/100 HEK cells observed in the presence of PMCA samples from the fifth cycle under the indicated conditions. **j**, The statistical analysis of the relative intensity of ProteoStat dye stained $\alpha S_{A53T}$-YFP aggregates in HEK cells treated with the PMCA sample from the fifth cycle in the absence and presence of SK-129 for 24 h. **k**, The statistical analysis of the relative viability of

HEK cells treated with the indicated conditions for 24 h determined using the MTT assay. The cell viability assays were conducted with at least four biological replicates and four technical replicates for each biological replicate and the reported error bars are the s.d.'s for every experiment. The counting of inclusions in HEK cells was carried out for six different experiments and for each experiment, 100 cells were counted from at least four different locations in an 8-well plate. For the ProteoStat dye staining of HEK cells, the experiment was conducted with at least four biological replicates and four technical replicates per biological replicate. The reported error bars are the s.d.'s for multiple set of experiments conducted on separate occasions. Statistical significance was analyzed using a one-way ANOVA with Tukey's multiple comparison's test. *$p<0.05$, **$p<0.01$, ***$p<0.001$.

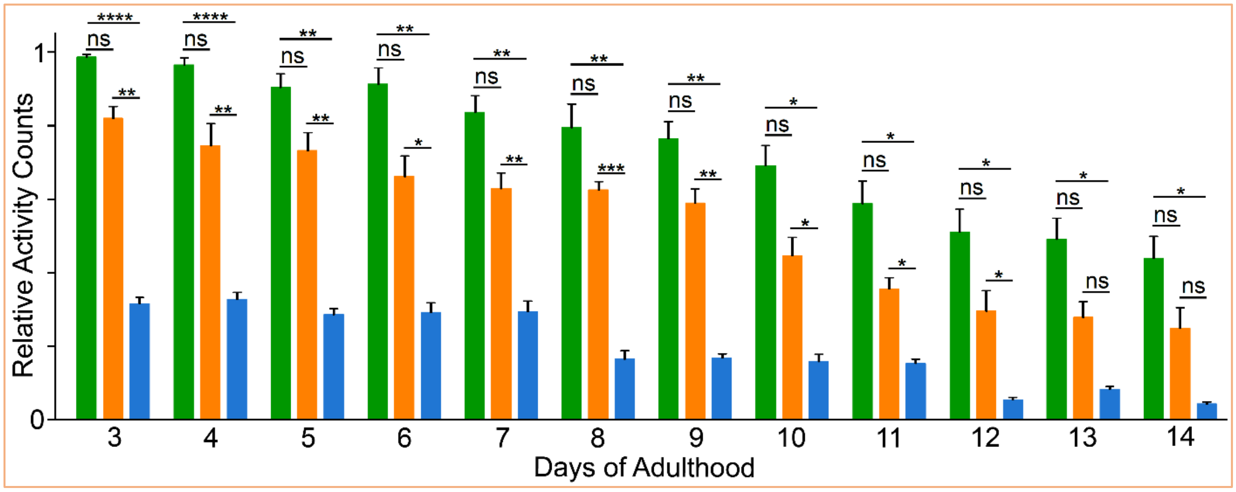


**Fig. 19.** The statistical analysis of the comparison of relative activity counts of the control strain (N2) and NL5901 in the absence and presence of 15 µM SK-129 was measured for 14 days of adulthood. The readings were taken for 1 h and for each reading, 100 worms were used for each condition. A total of 100 worms were used in duplicate for each experiment and each condition consisted of at least four independent experiments. The reported relative activity counts is an average of at least three separate experiments conducted in duplicate. The reported error bars are the s.d.'s for three sets of experiments conducted on separate occasions in duplicate. Statistical significance was analyzed using a one-way ANOVA followed by Tukey's multiple comparison test. *p<0.05, **p<0.01, ***p<0.001, ****p<0.0001.

### CITATIONS


1. Kumar, S., Birol, M., Schlamadinger, D.E., Wojcik, S.P., Rhoades, E. & Miranker, A.D. Foldamer-mediated manipulation of a pre-amyloid toxin. *Nat. Commun*. 7, 11412 (2016).
2. Kumar, S., Birol, M. & Miranker, A.D. Foldamer scaffolds suggest distinct structures are associated with alternative gains-of-function in a preamyloid toxin. *Chem. Commun*. 52, 6391-6394 (2016).
